# Chloroplasts Require Glutathione Reductase to Balance Reactive Oxygen Species and Maintain Efficient Photosynthesis

**DOI:** 10.1101/588442

**Authors:** Stefanie J. Müller-Schüssele, Ren Wang, Desirée D. Gütle, Jill Romer, Marta Rodriguez-Franco, Martin Scholz, Volker M. Lüth, Stanislav Kopriva, Peter Dörmann, Markus Schwarzländer, Ralf Reski, Michael Hippler, Andreas J. Meyer

**Affiliations:** Institute of Crop Science and Resource Conservation (INRES), University of Bonn, Friedrich-Ebert-Allee 144, 53113 Bonn, Germany; Institute of Plant Biology and Biotechnology, University of Münster, Schlossplatz 8, 48143 Münster, Germany; Plant Biotechnology, Faculty of Biology, University of Freiburg, Schänzlestr.1, 79104 Freiburg, Germany; Institute of Molecular Physiology and Biotechnology of Plants, University of Bonn, 53115 Bonn, Germany; Faculty of Biology, Cell Biology, University of Freiburg, Schänzlestr. 1, 79104 Freiburg, Germany; Botanical Institute, Cluster of Excellence on Plant Sciences (CEPLAS), University of Cologne, Cologne, Germany; Signalling Research Centres BIOSS and CIBSS, University of Freiburg, Schänzlestr.18, 79104 Freiburg, Germany

**Keywords:** Chloroplast, glutathione redox potential, light stress, *Physcomitrella patens*, redox-sensitive GFP, ROS

## Abstract

1. Thiol-based redox-regulation is vital to coordinate chloroplast functions depending on illumination. Yet, how the redox-cascades of the thioredoxin and glutathione redox machineries integrate metabolic regulation and reactive oxygen species (ROS) detoxification remains largely unresolved. We investigate if maintaining a highly reducing stromal glutathione redox potential (*E*_GSH_) via glutathione reductase (GR) is necessary for functional photosynthesis and plant growth.
2. Since absence of the plastid/mitochondrial GR is embryo-lethal in *Arabidopsis thaliana*, we used the model moss *Physcomitrella patens* to create knock-out lines. We dissect the role of GR in chloroplasts by *in vivo* monitoring stromal *E*_GSH_ dynamics, and reveal changes in protein abundances by metabolic labelling.
3. Whereas stromal *E*_GSH_ is highly reducing in wildtype and clearly responsive to light, the absence of GR leads to a partial oxidation, which is not rescued by light. Photosynthetic performance and plant growth are decreased with increasing light intensities, while ascorbate and zeaxanthin levels are elevated. An adjustment of chloroplast proteostasis is pinpointed by the induction of plastid protein repair and degradation machineries.
4. Our results indicate that the plastid thioredoxin and glutathione redox systems operate largely independently. They reveal a critical role of GR in maintaining efficient photosynthesis.

## Introduction

In photosynthetic eukaryotes, changes in environmental conditions, such as light intensity or temperature, provoke changes in electron flow both in the chloroplasts and in the mitochondria. Several mechanisms serve to rapidly modulate or redirect electron flow to minimise over-reduction of the two electron transport chains (ETCs), which can otherwise give rise to excessive formation of reactive oxygen species (ROS) (Schwarzländer & Finkemeier, 2013; Schöttler & Toth, 2014). Moreover, ROS serve as important signalling molecules in stress acclimation (Suzuki *et al*., 2012; Dietz *et al*., 2016). This implies that the rates of ROS generation and scavenging must be precisely balanced in these organelles. The maintenance of cellular redox pools for metabolism, antioxidant defence, and thiol-based redox switching requires the constant influx of electrons via light- and NADPH-powered redox cascades, involving the oxidation and reduction of cysteines in thioredoxins (Trx) and glutathione (Meyer *et al*., 2012; Yoshida & Hisabori, 2016; Geigenberger *et al*., 2017; Gütle *et al*., 2017).

Reduced glutathione (GSH) is present in cells at low millimolar concentrations (Meyer *et al*., 2001). Glutathione functions include ascorbate regeneration via the ascorbate-glutathione-cycle and detoxification of potentially toxic organic electrophils and heavy metals, as well as acting as a cofactor of monothiol glutaredoxins (Grx) for coordination of iron-sulfur clusters (Foyer & Noctor, 2011; Moseler *et al*., 2015). In addition, glutathione is also used as a substrate of dithiol Grx-catalysed protein (de)glutathionylation (Meyer *et al*., 2012; Zaffagnini *et al*., 2019). For the latter functions it is essential that glutathione can reversibly switch between its reduced form GSH and the oxidised form glutathione disulfide (GSSG) which involves the transfer of 2 electrons. The glutathione redox potential (*E*_GSH_) is dependent on the GSH concentration, as well as on the balance between GSH and GSSG. The *E*_GSH_ can vary drastically between subcellular compartments (Meyer, 2008; Kojer *et al*., 2012). In unstressed plant cells, the *E*_GSH_ of cytosol, peroxisomes, mitochondrial matrix and plastid stroma is highly reducing between −310 to −360 mV (Meyer *et al*., 2007; Schwarzländer *et al*., 2008). In these compartments, GSSG is efficiently regenerated to GSH by the action of glutathione reductase (GR) using NADPH as electron donor. In the cytosol and the mitochondria of *Arabidopsis thaliana* the loss of GR is partially compensated for by the presence of the NADPH-dependent Trx reductases A and B (NTRA, B) (Marty *et al*., 2009). Nevertheless, decreased GR activity of the plastid/mitochondria-targeted isoform leads to reduced root growth in seedlings (Yu *et al*., 2013). However, a complete loss of GR in plastids causes embryo-lethality (Marty *et al*., 2009; L. Marty & A.J. Meyer, unpublished).

Since this limits the usability of Arabidopsis as a model for GR function in green plastids, the significance of GR in stromal *E*_GSH_ maintenance, for photosynthesis and other plastid functions has remained unclear. Further, it is unknown to what extent the Trx system can serve as a back-up for the glutathione redox system in plastids. Here, we investigate the effects of the complete loss of plastid/mitochondria-localized GR in photosynthetic cells, utilising the moss *Physcomitrella patens* as a model (Reski, 2018). We assess growth and photosynthetic parameters, monitor the dynamics of plastid *E*_GSH_ by redox-sensitive GFP (roGFP)-based *in vivo* imaging and compare protein abundances between wildtype (WT) and GR mutants in response to a shift from low light to high light by quantitative proteomics.

## Material and Methods

### Plant materials and growth conditions

*Physcomitrella patens* (Hedw.) B.S. ecotype ‘Gransden 2004’ (International Moss Stock Centre (IMSC, http://www.moss-stock-center.org), accession number 40001) was grown axenically in agitated liquid Knop medium (250 mg l^−1^ KH_2_PO_4_, 250 mg l^−1^ KCl, 250 mg l^−1^ MgSO_4_ × 7H_2_O,1 g l^−1^ Ca(NO_3_)_2_ × 4H_2_O and 12.5 mg l FeSO_4_ × 7H_2_O, pH 5.8) (Reski & Abel, 1985) with micro-elements (ME) (H_3_BO_3_, MnSO_4_, ZnSO_4_, KI, Na_2_MoO_4_ × 2H_2_O, CuSO_4_, Co(NO_3_)_2_) (Egener *et al*., 2002) in a growth cabinet under long day conditions (16 h:8 h, light:dark, 22°C) at 100 µmol photons m^−2^ s^−1^. For phenotypic and pigment analyses, *P. patens* was grown on KNOP ME agar plates (12 g l^−1^ purified agar, Oxoid) at the indicated light intensity.

For measurements of photosynthetic parameters and preparation of proteins samples for MS/MS, *P. patens* protonema tissue was propagated under axenic conditions either on 9 cm or 4.5 cm Petri dishes overlaid with a cellophane disk on solidified PpNO_3_ medium (0.8 % (w/v) agar), or in glass flasks with PpNO_3_ medium (Gerotto *et al*., 2016) in a growth chamber under low light conditions (LL, 15 µmol photons m^−2^ s^−1^) at 25°C with a 16 h:8 h light:dark photoperiod. For control and high light assays, 10-day-old protonema plates were moved from LL to 50 µmol photons m^−2^ s^−1^ (CL) and 450 µmol photons m^−2^ s^−1^ (HL) respectively, maintaining temperature and photocycle.

### Generation of PpGR1 knock-out lines

A knock-out construct for Pp1s13_127V6.1 was designed by amplifying homologous regions from genomic DNA with primer pairs PpGR1ko_5PHR_F/ PpGR1ko_5PHR_R and PpGR1ko_3PHR_F/ PpGR1ko_3PHR_R (Table S1), introducing BspQ1 restriction sites to the homologous ends. DNA fragments containing homologous regions were joined with an expression cassette (Nopalin synthase promoter and terminator) for hygromycin phosphotransferase (hpt) via triple-template PCR. This construct eliminates a large part of the Pp1s13_127V6.1 coding sequence including the active site (Fig. S1a). The knock-out construct was subsequently ligated into the pJET1.2 (Thermo Scientific) vector, digested with BspQ1 and introduced via PEG-mediated protoplast transformation (Hohe *et al*., 2004) into a newly generated line expressing *TKTP-Grx1-roGFP2* (Schwarzländer *et al*., 2008; Speiser *et al*., 2018) stably integrated at the PTA2 locus under the control of the *PpActin5* promoter (Kubo *et al*., 2013; Mueller & Reski, 2015). In regenerated plants that survived hygromycin selection, integration of the construct into the target locus was verified using primer pairs spanning the 5’ integration site (5P_F and H3b_R, Fig. S1b) and the 3’ integration site (NosT_F and 3P_R) (Table S1). Absence of *PpGR1* transcript for independent knock-out lines was confirmed using the primer pair PpGR1_RT_F and PpGR1_RT_R in a reverse transcription PCR (Fig. S1c, Table S1). Moss lines are available from the IMSC under the accession numbers: TKTP-Grx1-roGFP2#40 IMSC 40836, Pp*gr1*#48 IMSC 40834, Pp*gr1*#88 IMSC 40835.

### Microscopy

Microscopy was carried out using a Zeiss LSM780 (attached to an Axio Observer.Z1) using a 25x (Plan-Apochromat 25x/0.8 Imm Korr NA0.8) or 40x (C-Apochromat 40×/1.2W Korr NA1.2) objective. Bright-field images were taken with an AxioCam MRc. Confocal laser scanning microscopy of roGFP2 redox state was achieved by consecutively exciting the roGFP2 with a 405 nm diode laser (at 2 % power output) and a 488 nm Argon laser (at 1 % power output) in line switching mode, using constant detector gain and emission from 508 to 535 nm. Autofluorescence was recorded after excitation at 405 nm and emission from 430 to 470 nm. Chlorophyll autofluorescence was monitored after 488 nm excitation at an emission of 680 to 735 nm. Image intensities and 405/488 nm ratios were calculated per pixel using a custom MATLAB-based software using background subtraction and autofluorescence correction (Fricker, 2016).

### Transmission electron microscopy

Transmission electron microscopy (TEM) was performed as described in Schuessele *et al*. (2016).

### NBT staining

Gametophores were stained in a 0.1 mg/ml nitro blue tetrazolium (NBT, Duchefa) solution in 75 mM potassium phosphate buffer (pH 7.0) for 1.5 h in the dark or in the light (120 µmol photons m^−2^s^−1^). Chlorophyll was subsequently removed by incubation in 80 % ethanol at 70°C (Lee *et al*., 2002).

### Ascorbate assay

Total and reduced ascorbate in *P. patens* samples was quantified according to Gillespie & Ainsworth (2007) with the modification that 2-2‘-bipyridyl was dissolved in 95 % ethanol. Five to 100 mg material was flash frozen in liquid nitrogen, homogenised in a bead mill (TissueLyser II, Qiagen, 30 Hz for 2x 1.5 min) and processed immediately.

### Pigment and glutathione analysis by HPLC

Photosynthetic pigments were quantified by high-performance liquid chromatography (HPLC) according to Thayer & Björkman (1990), see also Supplemental Information. Glutathione was extracted from c. 30 mg protonema tissue (grown in c. 100 µmol photons m^−2^s^−1^) in 10-fold volume of 0.1 M HCl. After centrifugation for 10 min at 4°C, 25 µL of the supernatant were neutralised with 25 µL 0.1 M NaOH and thiols reduced with 1 µL 0.1 M dithiothreitol for 15 min at 37°C in darkness. Ten μL 1 M Tris/HCl pH 8.0, 35 μL water were added and the GSH was derivatised using 5 μL of 0.1 M monobromobimane (Thiolyte^®^ MB, Calbiochem) in darkness for 15 min at 37°C. The reaction was stopped by adding 100 μL 9 % acetic acid and centrifugation at 4°C for 15 minutes. The bimane derivates were separated via HPLC (Spherisorb™ ODS2, 250 × 4.6 mm, 5 µm, Waters, Eschborn, Germany) using a linear gradient from 4 to 20 % of buffer A (90 % methanol (v/v), 0.25 % acetic acid (v/v), pH 3.9) in buffer B (10% methanol (v/v), 0.25 % acetic acid (v/v), pH 3.9) and detected fluorometrically with excitation at 390 nm and emission at 480 nm.

### Metabolic labelling and MS/MS analysis

For isotopic labelling of *P. patens*, Ca(NO_3_)_2_ × 4H_2_O in solid and liquid PpNO_3_ media was replaced by Ca(^15^NO_3_)_2_ × 4H_2_O (Cambridge Isotope Laboratories, England). To obtain fully labelled protonema tissue the protonema cultures were weekly sub-cultivated on fresh ^15^N labelled solid PpNO_3_ media for at least 4 months. To quantify differences in protein abundance between WT and *Δgr1* background, labelled and unlabelled protonema of cpGrx1roGFP2 #40 and *Δgr1* #48 were shifted from LL to HL for 1 h, including one label swap for each light intensity.

For protein extraction, c. 500 mg of *P. patens* protonema tissue was harvested and the surface water removed. The samples were frozen in liquid nitrogen and 3x homogenised using a MM300 mill (Retsch) for 30 s with a frequency of 30 s^−1^. Subsequently, 200-300 µl of protein extraction buffer (25 mM Trizma base, 1 % (w/v) SDS, 5 mM EDTA, 0.5 mM PMSF, 0.5 mM Benzamidine) were added, centrifuged at 2000 × g for 2 min and protein concentration determined in the supernatant by bicinchoninic acid assay. Equal protein amounts from ^14^N and ^15^N-labelled samples were mixed and further processed in a filter-aided sample preparation protocol for MS-based analysis according to Wiśniewski *et al*. (2009). All LC-MS/MS analyses were carried out on a system composed of an Ultimate 3000 RSLCnano UPLC coupled via a nanospray interface to a Q Exactive Plus mass spectrometer (Thermo Fisher Scientific).

### Shotgun Quantification

Peptides were pre-concentrated and desalted for 3 min on a trap column (Acclaim PepMap 100, 300 µM × 5 mm, 5 µm particle size, 100 Å pore size, Thermo Fisher Scientific) using 2 % (v/v) acetonitrile/0.05 % (v/v) trifluoroacetic acid in ultrapure water at a flow rate of 10 µl/min. Gradient separation of peptides was performed on a reversed phase column (C18, Acclaim Pepmap, 75 µm × 50 cm, 2 µm particle size, 100 Å pore size, Thermo Fisher Scientific) at a flow rate of 300 nl/min using the eluents 0.1 % (v/v) formic acid in ultrapure water (A) and 80 % (v/v) acetonitrile/0.1 % (v/v) formic acid in ultrapure water (B). The following gradient was applied: 2.5-18 % B (v/v) over 105 min, 18-32 % B (v/v) over 55 min, 32-99 % B (v/v) over 5 min, 99 % B (v/v) for 20 min.

MS full scans (MS1, *m/z* 300-1600) were acquired in positive ion mode at a resolution of 70,000 (FWHM, at m/z 200) with internal lock mass calibration on *m/z* 445.120025. For MS2 the 12 most intense ions were fragmented by higher-energy c-trap dissociation (HCD) at 27 % normalized collision energy (isolation window size: 1.5 m/z). Resolution for MS2 scans: 17,500 (FWHM, at *m/z* 200), target values for automatic gain control (AGC): 1×10^6^ and 5×10^4^ for MS full scans and MS2, respectively. The intensity threshold for MS2 was set to 1×10^4^. Maximum fill times were 50 ms (MS1) and 55 ms (MS2). Unassigned charge states, charged state 1 and ions with charge state 5 and higher were rejected.

### Bioinformatic analyses

LC-MS/MS data was processed with Proteome Discoverer (PD, version 2.2, Thermo Fisher Scientific). Raw files were searched using the SequestHT algorithm against a *P. patens* protein database based on the V1.6 gene models (Zimmer *et al*., 2013) supplemented with common contaminant proteins (cRAP, www.thegpm.org/crap/) with the following settings: Precursor and fragment mass tolerances 10 ppm and 0.02 Da, respectively; minimum peptide length: 6; maximum of missed cleavages: 2; variable modifications: Oxidation of methionine, N-acetylation of protein N-termini. For the identification of ^15^N-labelled peptides, a second database search was performed with ^14^N to ^15^N substitution(s) set as static modifications for all amino acids. Peptide-spectrum-matches (PSMs) were filtered using the Percolator node to satisfy a false discovery rate of 0.01 (based on q-values). Identifications were filtered to achieve a peptide and protein level FDR of 0.01. LC-MS/MS runs were chromatographically aligned with a maximum retention time drift of 10 min. Precursor ion quantification was performed using unique and razor peptides. Abundances were normalized to the maximum total peptide abundance in all files. Protein ratios (Δ*gr1* vs. WT) were calculated using the ‘pairwise ratio based’ approach with subsequent hypothesis testing (background based t-test) for the calculation of p-values.

The mass spectrometry proteomics data have been deposited to the ProteomeXchange Consortium (http://proteomecentral.proteomexchange.org) via the PRIDE partner repository (Vizcaíno *et al*., 2013) with the dataset identifier <PXD012843>.

### NPQ measurements

*In vivo* fluorescence in *P. patens* was measured with a Maxi-Imaging PAM chlorophyll fluorometer (Heinz Walz). Before measurements, LL, CL and HL grown protonema plates were dark-adapted for 40 min. NPQ (non-photochemical quenching) was calculated as (Fm–Fm’)/Fm’. Fv (the variable fluorescence) was calculated as Fv=Fm–Fo. The Fv/Fm ratio was used to evaluate the maximum PSII fluorescence in the fully dark-adapted state. Fm and Fm’ represent the maximum PSII fluorescence in the dark-adapted state and in any light-adapted state, respectively, and Fo represents the minimum PSII fluorescence in the dark-adapted state (Kukuczka *et al*., 2014).

### Spectroscopic measurements of photosynthetic parameters

Protonema from LL, CL and HL treated plates was measured with cellophane in buffer (Hepes 20 mM pH 7.5, KCl 10 mM). LEF+CEF (linear plus cyclic electron flow) and CEF (cyclic electron flow) were measured by following the relaxation kinetics of the carotenoid electrochromic band shift at 520 nm (corrected by subtracting the band shift at 546 nm) in the absence or presence of 10 μM DCMU and hydroxylamine, respectively. CEF and LEF+CEF were calculated as e^−1^s^−1^PSI^−1^ upon normalization to the PSI amount. The electrochromic shift signal upon excitation with a single saturating turnover flash (5 ns laser pulse) in the presence or absence of 10 μM DCMU and 1 mM hydroxylamine were used to estimate the PSI and PSI+PSII amount. DCMU and hydroxylamine in this measurement were used to fully block PSII photochemistry to facilitate the determination of the PSI amount (Terashima *et al*., 2012; Gerotto *et al*., 2016).

To measure the proton motive force (*pmf*), which consists of ∆pH (trans-thylakoid proton gradient) and membrane potential (∆ψ), 5-day-old protonema tissue from liquid cultures was harvested, dark-adapted for 15 min before analysis and exposed for 1:30 h to 300 µmol photons m^−2^ s^−1^ to obtain the steady-state of the ECS signal. Afterwards, a 1 min dark-phase was recorded (5 min of illumination in between measurements) to obtain at least 3 technical replicates at 520 nm and 546 nm respectively. g_H_^+^, which reflects the proton conductivity of the ATP synthase, was estimated by fitting the first 300 ms of the decay curve with a first-order exponential decay kinetic, ∆pH and ∆ψ were calculated as described previously (Wang *et al*., 2015).

## Results

### Moss lacking PpGR1 is viable and displays reduced growth

Loss of plastidic GR in Arabidopsis leads to embryo lethality (L. Marty & A.J. Meyer, unpublished), indicating an essential role in the non-photosynthetic tissues of early sporophyte development, but also preventing further studies of GR function in green tissues. We hence chose *Physcomitrella patens* as a model because knock-out mutants can be generated using protoplastation and regeneration of photosynthetically active vegetative cells, circumventing embryogenesis and non-photosynthetic tissue (e.g. Schween *et al*. (2005)). We hypothesized that null mutants of organellar GR might be viable in plants that maintain green plastids throughout their life cycle. We therefore generated knock-out constructs replacing exons two to five containing the translation start site and the active site in the gene encoding the previously identified dual-targeted mitochondria and plastid-localised glutathione reductase Pp1s13_127V6.1 (named *GR1* in *P. patens*, Xu *et al*., (2013)) with a hygromycin resistance cassette via homologous recombination (Fig. S1a). As genetic background, we utilised a newly generated *P. patens* line expressing the plastid-targeted *E*_GSH_ biosensor Grx1-roGFP2 under the control of the *P. patens Actin 5* promoter (Weise *et al*., 2006; Mueller & Reski, 2015). We isolated plants surviving hygromycin selection under constant light. They were genotyped for the integration of the knock-out construct at the target locus (Fig. S1b), and the absence of *PpGR1* transcript was confirmed (Fig. S1c). All plants lacking *GR1* transcript showed a dwarf phenotype (Fig. 1a) compared to the WT line and the line expressing plastid-targeted Grx1-roGFP2 (cpGrx1roGFP2 #40, Fig.1a). Two independent lines, *Δgr1* #48 and *Δgr1* #88, were chosen for further analysis. As the *Arabidopsis thaliana gr2-1* null mutant is embryo-lethal, we tested if *Δgr1* knock-out mutants were able to complete the moss life cycle. Under inducing conditions, *Δgr1* #48 and *Δgr1* #88 formed sporophytes that underwent complete development and opened to release mature spores (Fig. 1b). Thus, GR1 is not necessary for embryo development in *P. patens*. However, spore germination of *Δgr1* #48 and Δgr1 #88 was delayed by several days, with spores being able to germinate eventually (Fig. 1b). Apart from the dwarfed appearance, the mutant lines grew fewer caulonema filaments and rhizoids than the WT (Fig. S2).

**Fig. 1:**
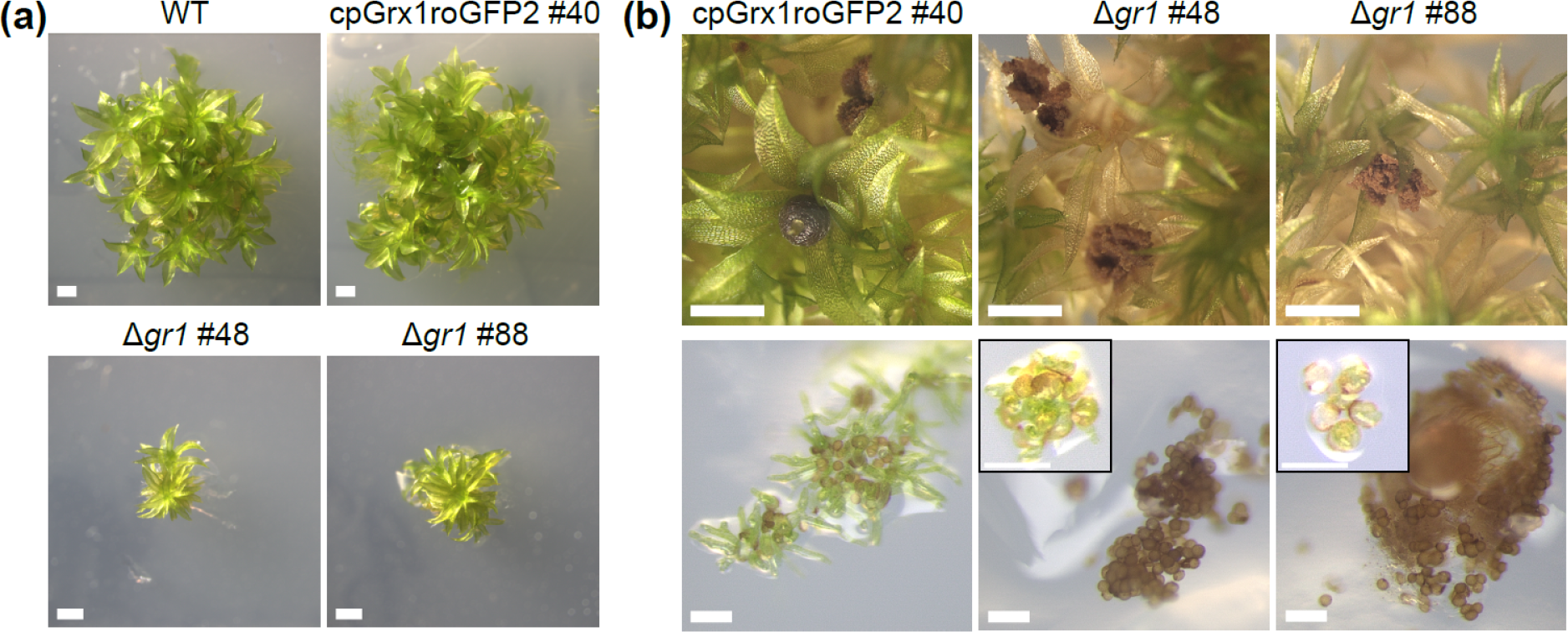
Phenotype of *Physcomitrella patens* plastid/mitochondrial glutathione reductase knock-out lines. (a). Wildtype (WT), a sensor line with expression of chloroplast-targeted Grx1-roGFP2 (cpGrx1roGFP2 #40) and two independent Δ*gr1* knock-out lines (Δ*gr1* #48, Δ*gr1* #88) after 40 d of growth on agar plates. Bars, 1 mm. (b) Gametophores carrying ripe sporophytes (upper panel) and spore germination after 8 d and 36 d (inlays Δ*gr1*). Bars,1 mm (upper panels), 0.1 mm (lower panels).

The ultrastructure of chloroplasts lacking GR1 was investigated using transmission electron microscopy (TEM), which revealed normally packed grana stacks and stroma lamellae, undistinguishable from WT (Fig. S3).

### Plastid redox state is dynamic in WT, shifted to less reducing values in *Δgr1* plants and not rescued via Trx reduction under light

Taking advantage of the stromal-targeted Grx1-roGFP2, the steady state of the chloroplast *E*_GSH_ was determined by confocal *in vivo* imaging of roGFP2 redox state (Fig. 2a). The fluorescence excitation ratio 405/488 nm ratio increases with sensor oxidation. It was 0.91 +/− 0.17 in WT, 1.61 +/− 0.25 in *Δgr1*# 48 and 1.81 +/− 0.28 in *Δgr1* #88 (Fig. 2a,b), indicating that the stromal *E*_GSH_ is less reducing in *Δgr1* lines. To test for stability of stromal *E*_GSH_ after exogenous reduction, plants were incubated with 2 mM dithiothreitol (DTT) and then exposed to continuous laser scanning under the confocal microscope. As a result, plastid Grx1-roGFP2 was rapidly re-oxidised in the *Δgr1* plants, but not in the WT background (Fig. 2c). This result prompted us to investigate the dynamics of *E*_GSH_ in dark-to-light and light-to-dark transitions (Fig. 2d). Dark-adapted plants were exposed to a dark-to-light transition (100 µmol photons m^−2^ s^−1^ for 10 min) during confocal imaging, resulting in a transient oxidation, followed by a reduction of the Grx1-roGFP2 redox state. The following light-to-dark transition (after 10 min of light exposure) resulted in an oxidation. Pre-incubation of plants with the electron transport inhibitor 10 µM DCMU (3-(3,4-dichlorophenyl)-1,1-dimethylurea) blocked the *E*_GSH_ dynamics, indicating a dependence on the photosynthetic ETC. No reduction of Grx1-roGFP2 in the light was observed in *Δgr1* plants. As Trx systems constitute a functional backup for cytosolic and mitochondrial GRs (Marty *et al*., 2009; L. Marty & A.J. Meyer, unpublished), we next investigated whether the survival of *Δgr1* plants was dependent on the activity of the ferredoxin-thioredoxin reductase (FTR) system. As FTR is supplied with electrons from the chloroplast ETC, we tested survival of the *Δgr1* mutant lines after transfer from constant light to short day conditions (Fig. S4a), and under extended darkness (44 d, Fig. S4b). These experiments revealed that *Δgr1* #48 and *Δgr1* #88 were not sensitive to incubation in darkness, suggesting that the FTR-dependent redox cascades are not required for the survival of the *P. patens* mutants.

**Fig. 2:**
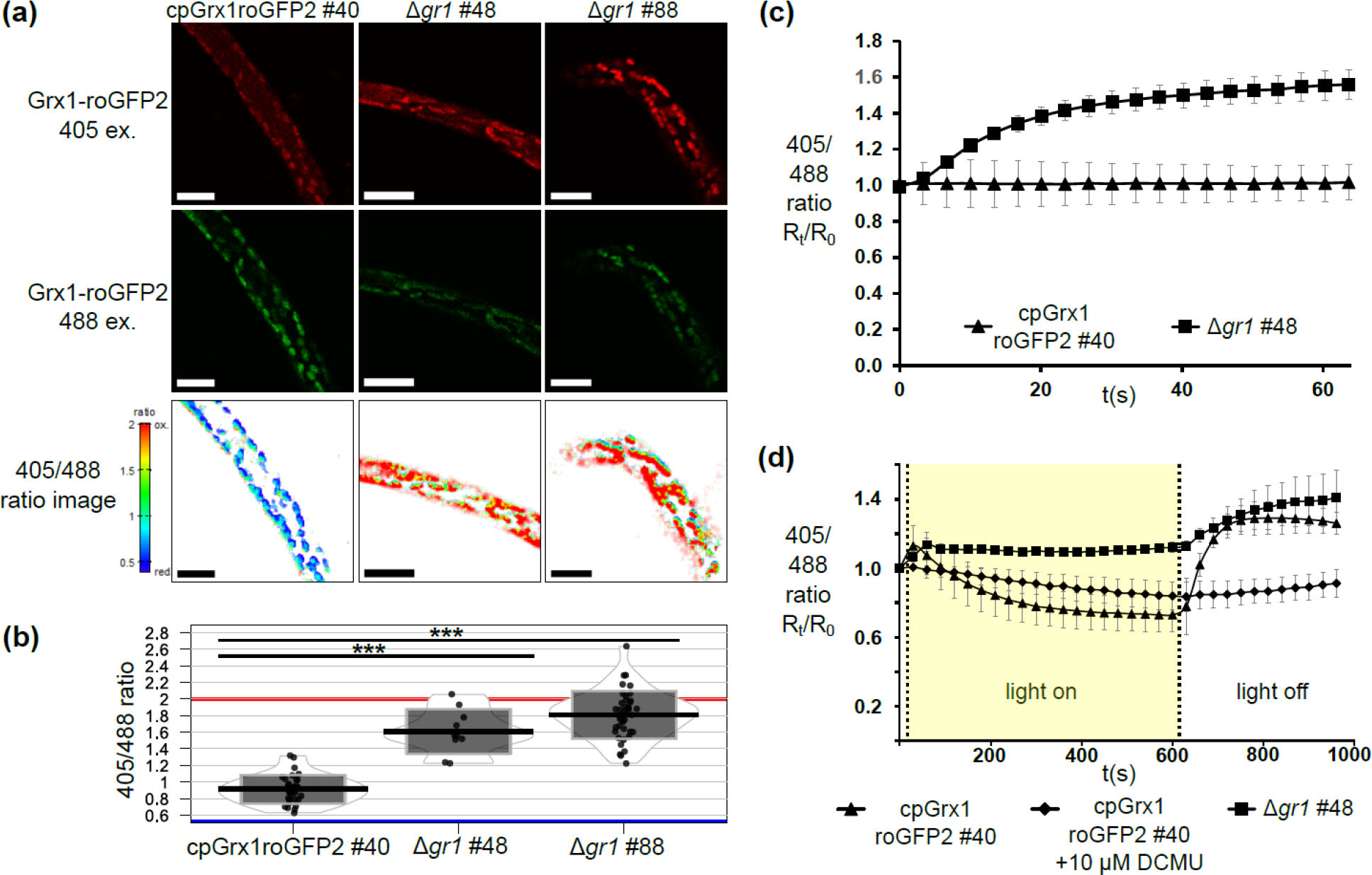
Chloroplast redox homeostasis in Δ*gr1* plants measured by Grx1-roGFP2. (a) The redox-state of chloroplast-targeted Grx1-roGFP2 was measured ratiometrically by excitation at 405 nm (upper panel, red), and 488 nm (middle panel, green). The emission ratio (lower panel) indicates a highly reduced roGFP2 in WT plastids whereas roGFP2 is nearly fully oxidised in Δ*gr1* lines. Bars, 20 µm. (b) 405/488 nm ratio of plastid Grx1-roGFP2 (*n*=10-30); plot depicts mean +/-SD as boxes, individual data points, and data point density. *** significant difference, *P*<0.001 (one-way ANOVA, TukeyHSD post hoc test). *In vivo* sensor calibration: Red line: 405/488 nm ratio of fully oxidised Grx1-roGFP2 (5 mM 2,2’-dipyridyldisulfide (DPS)); blue line: 405/488 nm ratio of fully reduced Grx1-roGFP2 (10 mM dithiothreitol (DTT)). (c) WT and Δ*gr1* mutant were pre-treated with 2 mM DTT to achieve Grx1-roGFP2 reduction and then exposed to continuous laser scanning in water (1 scan/timepoint; WT background *n*=4, *Δgr1 n*=9). (d) *In vivo* measurement of light-dependent chloroplast *E*_GSH_ dynamics. Dark-adapted WT (cpGrx1roGFP2 #40) and Δ*gr1* plants were exposed to a dark/light/dark transition (100 µmol photons m^−2^ s^−1^ for 10 min). Incubation with the electron transport inhibitor 10 µM DCMU blocked the observed *E*_GSH_ dynamics; *n*=3.

### *Δgr1* plants show altered responses of reactive oxygen species dynamics and photosynthetic function

As *Δgr1* knock-out plants were not sensitive to dark-incubation (Fig. S4) and showed an oxidative response to the laser light used for microscopic imaging (Fig. 2c), their growth habit under different light intensities was tested (Fig. S5a). Mutants did not profit from increasing light fluencies (30 to 130 µmol photons m^−2^ s^−1^), as their fresh weight did not increase, in contrast to WT (Fig. S5b). In order to investigate the impact of different light intensities on ROS scavenging, the levels of ascorbate and dehydroascorbate were determined. The measurements revealed a higher total level of ascorbate in *Δgr1* #48 and *Δgr1* #88 that reached 500 % to 700 % of WT levels in different light intensities (Fig. 3a). Interestingly, dehydroascorbate levels were increased as well, but were present at the same ratio to reduced ascorbate as in WT (on average 8.1+/-2.5 % of total ascorbate, Fig. 3b). We incubated gametophores with nitro blue tetrazolium (NBT) in the dark or in light as a means to detect superoxide (O_2_^•-^), and found increased staining intensity in light-incubated *Δgr1* plants (Fig. 3c). To induce additional ROS formation by photosystem I (PSI), single gametophores were transferred to medium containing 1 µM Paraquat. While WT plants were still able to grow, *Δgr1* #48 and *Δgr1* #88 plants bleached and died (Fig. 3d, upper panel). To stop photosynthetic electron flow, moss colonies were transferred to photoheterotrophic growth conditions and their survival on DCMU was tested according to Bricker *et al*. (2014). WT plants survived when sucrose (1% (w/v)) was present in the medium, whereas *Δgr1* #48 and *Δgr1* #88 plants bleached and died (Fig. 3d, middle panel).

**Fig. 3:**
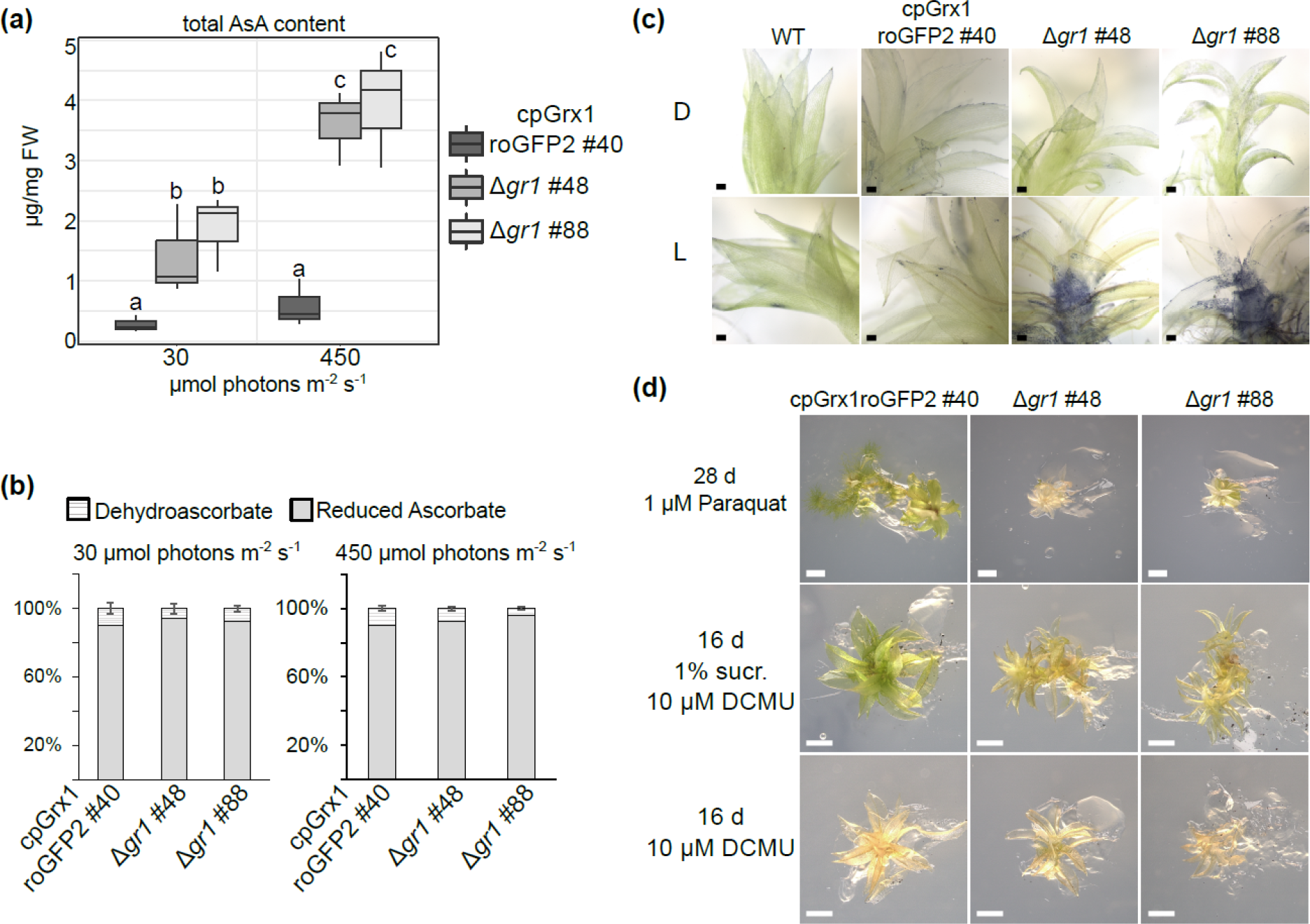
Ascorbate levels and resistance reactive oxygen species. (a) Total ascorbate (AsA) content of a WT line with the Grx1-roGFP2 sensor and two Δ*gr1* lines under different light conditions. Lower case letters indicate significant differences (*P*<0.05, two-way ANOVA, Tukey HSD post hoc test). (b) Relative contents of dehydroascorbate and reduced ascorbate in WT and Δ*gr1* plants under different light conditions. (c) Nitro blue tetrazolium (NBT) staining of moss gametophores either kept for 1.5 h in the dark (D) or in the light (120 µmol photons m^−2^ s^−1^, L). Bars, 100 µm. (d) Plant growth in the presence of Paraquat (upper panel). Survival after transfer to photoheterotrophic growth conditions in the presence of DCMU (3-(3,4-dichlorophenyl)-1,1-dimethylurea) (middle panel) and control of lethal effect of DCMU under photoautotrophic conditions (lower panel). Representative images (*n*=6-9) shown. Bars, 1 mm.

Under exposure to higher light fluencies (HL, 450 µmol photons m^−2^ s^−1^), white sectors appeared in leaflets of *Δgr1* plants (Fig. 4a). When HL was additionally combined with elevated temperature *Δgr1* #48 and *Δgr1* #88 plants died, whereas WT and *Δgr1* mutants were able to recover from temperature stress only (Fig. 4a). An examination of photosynthetic parameters revealed a light intensity-dependent decrease of non-photochemical quenching (NPQ), linear and cyclic photosynthetic electron flow (LEF+CEF) and the photosystem I to photosystem II ratio (PSI/PSII) (Fig. 4b). While the NPQ amplitude decreased with increasing light intensity in *Δgr1*, the NPQ dark relaxation was slower under all light conditions tested. CEF was not increased under different light intensities, but showed a slight increase after 6 h induction via anoxia (Fig. S6a,b). The Fv/Fm ratio decreased in *Δgr1* with increasing light, indicating decreased photosynthetic efficiency (Fig. S6a). At the same time, total chlorophyll (a+b) levels were decreased (Fig. S6c).

**Fig. 4:**
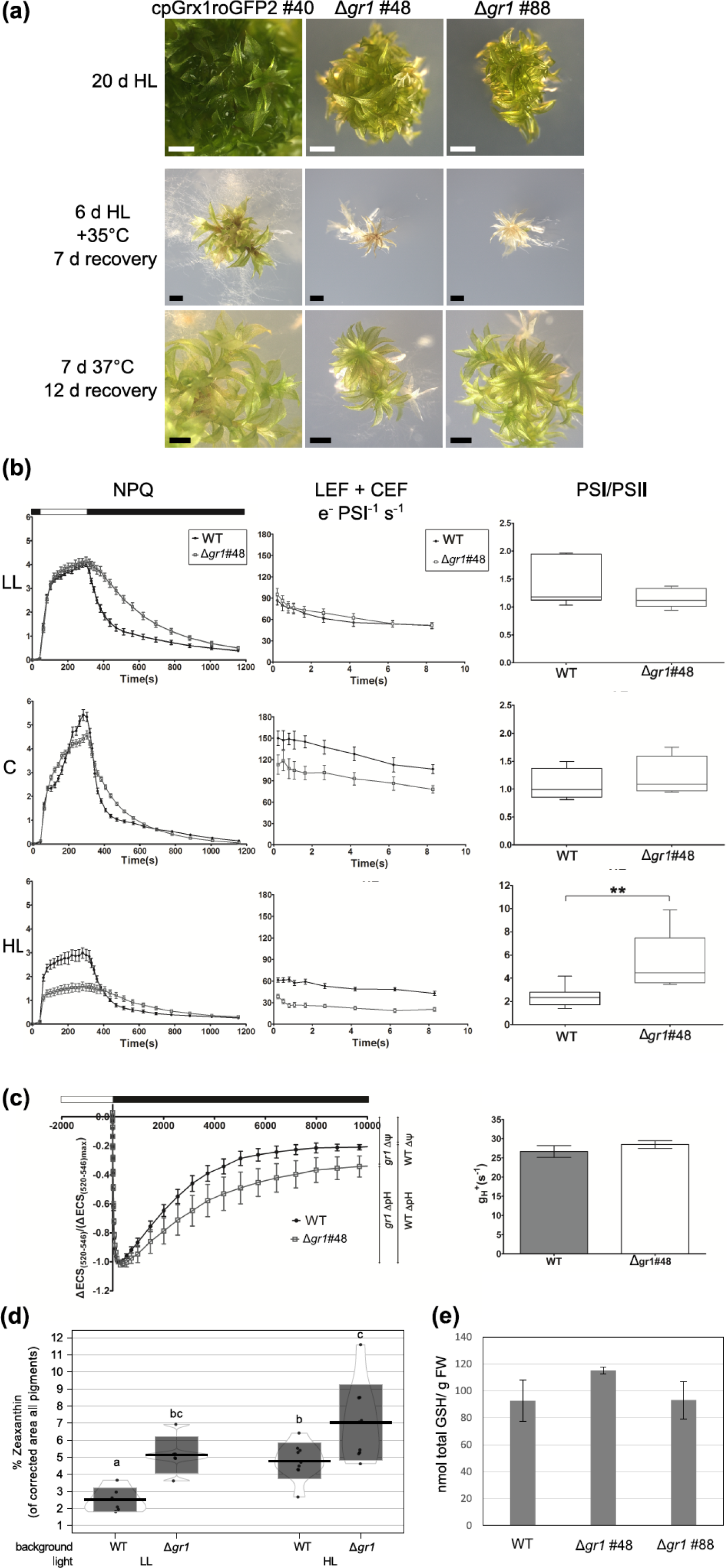
Light-sensitivity of Δ*gr1* plants. (a) High light treatment of plants grown on agar plates (HL, upper panel); bars, 0.5 mm. High light combined with elevated temperature (middle panel), and elevated temperature in the dark (lower panel). Bars, 1 mm. (b) Photosynthetic parameters measured under low light (LL, 15 µmol photons m^−2^ s^− 1^), control light (C, 50 µmol photons m^−2^ s^−1^) and high light (HL, 450 µmol photons m^−2^ s^−1^). Non-photochemical quenching (NPQ, left panel). Bars indicate standard deviation (SD, *n*=6); white and black boxes on top indicates light and dark phase of the measurements. Actinic light (1076 μmol photons m^−2^⋅s^−1^) was switched off after 5 min illumination. Photosynthetic linear and cyclic electron flow (LEF+CEF; middle panel; *n*=6 LL, CL; *n*=12 HL). PSI/PSII ratio (right panel; *n*=6 LL, CL; *n*=12 HL). (c) Measurement of the proton motive force components pH gradient and proton conductivity of the ATP synthase (*n*=3). Left: Dark relaxation of the carotenoid electrochromic shift signal (ECS) after illumination (white box) with 300 μmol photons m^−2^⋅s^−1^ (*P*<0.0001, paired t-test). White and black box on top indicates light and dark phases of the measurements. Membrane potential ΔΨ. Right: Proton conductivity of the ATP synthase g_H_^+^. (d) HPLC measurements of zeaxanthin levels in plants grown under low light (LL, 15 µmol photons m^−2^ s^−1^) and plants grown under control light (C, 50 µmol photons m^−2^ s^−1^) and shifted for 4 h to high light (HL, 450 µmol photons m^−2^ s^−1^). *n*=6, small letters depict significant differences, *P*<0.05 (two-way ANOVA, Tukey HSD post hoc test), plot depicts mean +/-SD as boxes, individual data points, and data point density. (e) HPLC measurements of total glutathione level (GSH+GSSG). No significant differences (one-way ANOVA, Tukey HSD post hoc test).

To study the proton motive force (*pmf*) in mutant chloroplasts in comparison to WT, electrochromic shift assays (Kramer & Crofts, 1989) were conducted and revealed a decreased pH gradient and increased membrane potential ΔΨ in *Δgr1*, resulting in a similar *pmf*. The proton conductivity of the ATP synthase was not significantly altered (Fig. 4c).

As NPQ relaxation was slower under all light conditions tested, similar to Arabidopsis *npq2* (zeaxanthin epoxidase) mutants (Niyogi *et al*., 1998), we measured levels of photosynthesis pigments and found that zeaxanthin levels were increased up to 2-fold relative to total pigment content in the *Δgr1* mutant in comparison to WT (Fig. 4d). Total glutathione levels (GSH+GSSG) were not significantly different to WT (Fig. 4e).

### Quantitative proteomics reveals light intensity-dependent protein level changes in Δ*gr1* plants

To assess the significance of GR function for proteostasis under changing light intensities we used metabolic labelling with the stable isotope ^15^N in combination with quantitative proteomics. Changes in protein abundances in WT and *Δgr1* plants upon a shift from low light (LL) to high light (HL) were investigated using quantitative proteomics (Fig. S7). The sensor expressing line cpGrx1roGFP2 #40 (WT) and Δ*gr1#48* were labelled *in vivo* by growth on medium containing Ca(^15^NO_3_)_2_ as exclusive nitrogen source, or non ^15^N-containing medium, respectively. Labelled and unlabelled samples of both lines were exposed to HL and samples taken after 1 h whereas control samples were kept at LL. For the same experimental condition, protein extracts from labelled and unlabelled samples of Δ*gr1* and cpGrx1roGFP2 #40 (WT), respectively, were mixed in a 1:1 ratio, tryptically digested and analysed by LC-MS/MS. To exclude any bias introduced by the labelling, one label swap experiment was performed per light intensity, resulting in a total of four experiments (2x Δ*gr1* vs. WT HL, 2x Δ*gr1* vs. WT LL, Fig. S7). In total, 1657 proteins with Δ*gr1* vs. WT ratios were quantified, of which 125 were differentially abundant (*P*<0.05) in Δ*gr1* and WT in LL or HL (Fig. 5a, blue dots). Of these 125 proteins, 70 were down-regulated and 59 up-regulated (four were either up- or downregulated, depending on the light conditions). The overlap between differentially regulated proteins in LL in comparison to HL was 8.6 % for the down-regulated proteins and 20.3 % for the up-regulated proteins (Fig. 5b). Following a manual annotation of subcellular localisation (Table S2), based on available organelle proteomics data sets for *P. patens* (Mueller *et al*., 2014) and SUBAcon (Hooper *et al*., 2014) annotation of Arabidopsis homologs, the largest fraction of regulated proteins was attributed to plastids (38), followed by the cytosol (20) and proteins with unclear localisation (13) (Fig. 5c). In addition, differentially regulated proteins were sorted into 29 functional categories (Table S2) and categories containing more than two proteins plotted to visualise category-specific down- or up-regulation in the different light conditions (Fig. 5d). Here, more proteins with unknown function were down-regulated specifically in LL or HL whereas several proteins with unknown function were up-regulated in both light conditions. In the categories “protein homeostasis” and “photosynthesis light reactions” more proteins were down-regulated in LL, but more proteins were up-regulated in HL, indicating a strong influence of the light intensity on protein levels in these categories. PSI subunit E (Pp1s101_2V6.1, 2^−0.33^) was less abundant in LL, whereas the plastid-encoded PSII subunits D1 and D2 were less abundant after the shift to HL (PsbA PhpapaCp046, 2^−0.25^; PsbD PhpapaCp044, 2^−0.2^). In the functional category “protein homeostasis”, under HL, the increase of one isoform of the plastid proteasome proteolytic subunit ClpP (Pp1s161_14V6.1, 2^6.64^) and of chaperonin 60 (Chp 60) alpha and beta subunits (Pp1s16_322V6.1 2^0.48^; Pp1s14_298V6.1, 2^0.47^; Pp1s15_485V6.1, 2^0.55^) indicated an increased demand for protein stabilization and degradation. Proteins of cytosolic translation were down-regulated under LL, whereas proteins of respiratory complex I and photosynthetic dark reactions were only affected in HL. Proteins of plastid translation were up-regulated in LL.

**Fig. 5:**
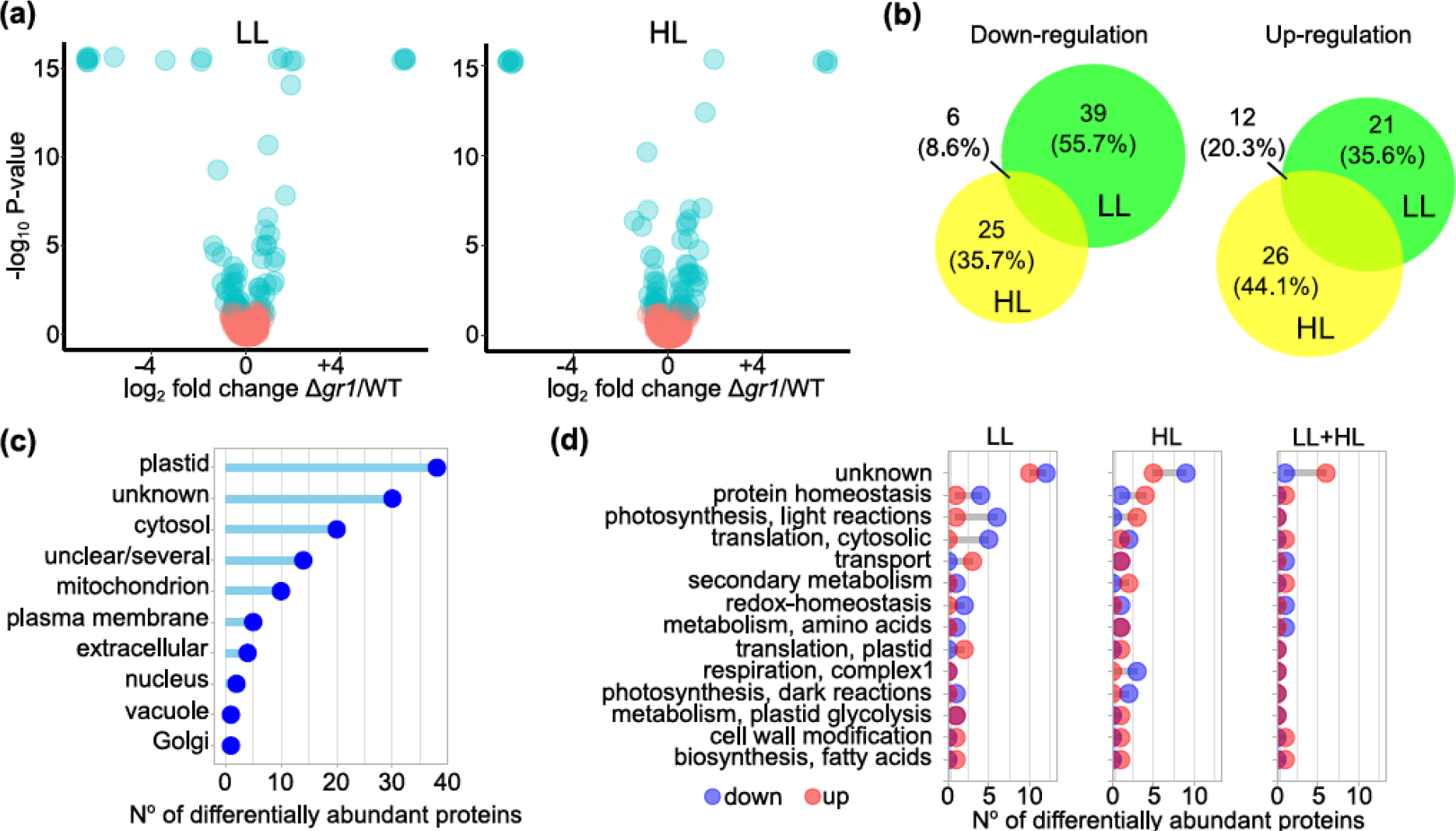
Quantification of protein abundances by metabolic labelling. (a) Protein abundances of Δ*gr1*/WT in low light (LL) and after a shift to high light (HL) for 1 h. (b) Area-proportional Venn diagram (Hulsen *et al*., 2008) showing overlap of proteins with differential abundance (Δ*gr1*/WT) between light treatments. (c) Comparison of manually annotated subcellular localisations (Table S2) of proteins with differential abundance. (d) Cleveland dot plot showing functional categories (Table S2) containing >2 proteins with differential abundance (Δ*gr1*/WT). LL, low light; HL, high light; LL+HL, differentially abundant in low light and high light.

Prominent changes in protein abundances include a plastid ribosome release factor (Pp1s130_293V6.1, 2^6.64^, HL) and a KEA (K^+^-efflux antiporter) homolog (Pp1s2_217V6.1, 2^−6.64^, HL) with high similarity to AtKEA1 and AtKEA2 (Kunz *et al*., 2014). Notably some enzyme isoforms were regulated differentially, such as enolase (Pp1s1_527V6.1, c. 2^1.2^, LL+HL; Pp1s37_237V6.1, 2^−6.64^, LL+HL).

As PpGR1 (Pp1s13_127V6.1) was also identified in the proteomics analysis as differentially abundant (Table S2), we confirmed the absence of PpGR1 additionally by a targeted proteomics approach (Fig. S8).

Further, we screened the dataset for known redox-regulated proteins and found one of the three isoforms of gamma subunit of chloroplast ATP synthase (Pp1s35_234V6.1, 2^−1.35^, LL), a putative plastid glucose-6-phosphate dehydrogenase (Pp1s338_65V6.1, 2^−0.36^, HL), as well as one of two FBPase isoforms (FBPase 2 Pp1s20_373V6.1 2^−0.87^, HL) down-regulated. Interestingly, *P. patens* possesses a putative oxidoreductase using GSSG with similarity to bacterial YfcG (Pp1s339_37V6.1) that was down-regulated under both light conditions.

## Discussion

### Stromal *E*_GSH_ responds to photosynthetic status

We generated viable null mutants of *P. patens* GR1, and found that absence of GR1 leads to a shift in the stromal *E*_GSH_. After Grx1-roGFP2 calibration, sensor 405/488 nm excitation ratio measurements can be translated into degree of sensor oxidation. As the redox potential of roGFP2 equilibrates with the redox potential of glutathione, *E*_GSH_ can be calculated, with the limitation that compartment pH has to be estimated (Meyer *et al*., 2007; Schwarzländer *et al*., 2008). In *Δgr1* plants, the degree of oxidation of the plastid-targeted *E*_GSH_ sensor Grx1-roGFP2 was severely shifted. Based on the change of the 405/488 nm ratio and the *in vivo* sensor calibration (Fig. 2b), we calculated a shift from c. 48 % oxidation in the WT background to c. 92 % oxidation in the Δ*gr1* lines. This would correspond to a 33 mV shift in the redox potential (calculated for pH8: −311 mV in WT vs. −278 mV in Δ*gr1*). In comparison, the redox potential in Arabidopsis epidermal plastids was determined as c. −361 mV at pH 8 (Schwarzländer *et al*., 2008). A shift of 30 mV would mean an increase of the relative amount of GSSG from 0.01 % to 0.1 %, calculated for a total concentration of 2.5 mM GSH (Meyer *et al*., 2007).

As in the same compartments glutathione- and Trx-dependent thiol switching fuelled by distinct reductases co-exists (Buchanan & Balmer, 2005), the cross-talk of these systems has been dissected for the cytosol and the mitochondria. Thus, cytosolic Trx redox-state can be rescued via the glutathione system (Reichheld *et al*., 2007). Vice versa, the NTRA/B system present in the cytosol and the mitochondria constitutes a functional backup of the glutathione system (Marty *et al*., 2009). However, reduction of many plastid Trxs is light-dependent and *E*_GSH_ in other cellular compartments is stable. It was hypothesized that the Trx-dependent redox cascades in plastids cannot provide sufficient backup to reduce the plastid *E*_GSH_ (L. Marty & A.J. Meyer, unpublished). Our data indicate that a shift of *E*_GSH_ in plastids occurs in the absence of GR, but that this shift is limited. The resulting steady state level may be a consequence of either electron flux to GSSG from Trxs, export of GSSG (Morgan *et al*., 2013; Noctor *et al*., 2013) from plastids or increased GSH biosynthesis (Choudhury *et al*., 2018). However, we did not find increased glutathione levels in the absence of GR1 (Fig. 4e). In dynamic measurements in ectopically reduced Δ*gr1* plants, the shifted stromal *E*_GSH_ was rapidly re-established by exposure to laser light, suggesting that GSSG rapidly accumulates in mutants upon illumination. As the regeneration of GSSG via GR is lacking in the mutant plastids, this likely represents GSSG formed by the ascorbate-GSH cycle, i.e. dehydroascorbate reductase (DHAR) activity. While the relative contributions of monodehydroascorbate reductase (MDHAR) and DHAR to the plastid ascorbate regeneration were debated (Asada, 1999; Polle, 2001), plastid-targeted AtDHAR3 was shown to contribute to ascorbate recycling with mutants being sensitive to high light (Noshi *et al*., 2016). The increase in total ascorbate and dehydroascorbate levels in Δ*gr1* mutants indicates impaired function of the ascorbate-GSH cycle consistent with a substantial contribution of plastid GR, including non-stress conditions. Light was not necessary for the survival of Δ*gr1* plants, indicating that light-dependent reduction of Trxs was not a prerequisite for viability. Further, cross-talk between the Trx- and glutathione redox cascades in plastids is limited, which is in contrast to previous findings in the cytosol and mitochondria (Reichheld *et al*., 2007; Marty *et al*., 2009).

Notably, in WT plants exposed to a successive dark/light and light/dark transition, the stromal *E*_GSH_ responded dynamically, showing that *E*_GSH_ is rapidly light-responsive in the presence of GR. In addition, after the transition from light to darkness, we observed a rapid rise of the stromal *E*_GSH_ pointing to oxidative processes in consequence of a light/dark transition. This oxidation is analogous to Trx oxidation that is required to deactivate redox-regulated Calvin Benson cycle enzymes (Wolosiuk & Buchanan, 1977; Yoshida *et al*., 2018). Peroxiredoxins have been reported to act as possible electron sinks (Pérez-Ruiz *et al*., 2017; Vaseghi *et al*., 2018).

At the measured stromal *E*_GSH_ in Δ*gr1*, still most of the stromal glutathione is present in the reduced state (99.9 %). Nevertheless, a shifted *E*_GSH_ is likely to affect downstream redox-cascades, as well as GSH-dependent enzymatic reactions. This includes glutaredoxins (Grx) and dehydroascorbate reductase (DHAR). While the involvement of plastid Grxs in iron-sulfur cluster coordination and protein (de)glutathionylation has been shown (Zaffagnini *et al*., 2012; Moseler *et al*., 2015; Rey *et al*., 2017; Zannini *et al*., 2019), only very few target proteins are currently known. Thus, the future challenge is to identify specific target cysteines affected by a shifting *E*_GSH_ in order to appraise its potential physiological role in redox regulation and signalling.

### Consequences of a lack in plastid/mitochondrial GR

The absence of plastid/mitochondrial GR resulted in a pronounced dwarfism as well as in light sensitivity of the mutant plants. As dual targeting of one GR isoform is evolutionarily conserved (Xu *et al*., 2013), the contribution of the lack of mitochondrial GR to the overall phenotype of the mutants is not resolved yet. However, in Arabidopsis it is the lack of GR in plastids, and not in mitochondria, that results in embryo-lethality (L. Marty & A.J. Meyer, unpublished; Nietzel *et al*., 2019). We found that in *P. patens*, the plastid/mitochondrial isoform of GR is not necessary for embryo development. This suggests that either (1) the process that causes embryo-lethality in Arabidopsis is not important for *P. patens* embryo development, or (2) that the lack of GR is compensated for in *P. patens* allowing embryo development to proceed. For instance, in contrast to flowering plant *gr* mutants under stress (Ding *et al*., 2012), *P. patens Δgr1* mutants showed a high increase in total ascorbate levels.

In the green haploid moss gametophyte, lack of GR1 caused slow growth as well as defects in photosynthetic parameters. Whereas photosynthetic electron flow was not affected in low light, increasing light fluencies resulted in decreased electron flow in photosynthetic light reactions, as well as a near complete loss of NPQ. In addition, the retarded relaxation of NPQ kinetics and increased zeaxanthin levels, similar to the Arabidopsis *npq2* mutant lacking zeaxanthin epoxidase (Niyogi *et al*., 1998) suggest an influence of *E*_GSH_ in the regulation of zeaxanthin epoxidase, also under non-stress conditions. Under HL, violaxanthin is converted into zeaxanthin which is involved in scavenging ROS as inferred from the analysis of Arabidopsis *vte1* mutant deficient in the synthesis of tocopherol, one of the lipid antioxidants in chloroplasts (Havaux *et al*., 2005). Violaxanthin de-epoxidase, the enzyme involved in zeaxanthin production, is activated by acidification of the lumen pH and uses ascorbate as cosubstrate (Arnoux *et al*., 2009).

Further, an increasing PSI/PSII ratio under higher light fluencies indicated PSII damage. Concomitantly, we found decreased levels of photosystem II subunits D1/D2 and increased protein levels of plastid chaperones and protein degradation, confirming an increased demand for protein repair and degradation in HL. PSII efficiency was already linked to GR activity in a study using a tobacco line with 30 % of plastid/mitochondrial GR activity (Ding *et al*., 2009). Here, the diminished GR activity resulted in decreased chlorophyll, ascorbate, DHA levels and PSII efficiency, as well as increased H_2_O_2_ levels under chilling stress (Ding *et al*., 2012). Concomitantly, overexpression of plastid/mitochondrial GR was beneficial under photoinhibitory conditions in poplar and cotton (Foyer *et al*., 1995; Kornyeyev *et al*., 2003).

In Δ*gr1* plants, we found an elevated ∆ψ and decreased ∆pH, resulting in a similar *pmf* and H^+^ conductivity (g_H_^+^) compared to WT. An elevated electric field component can increase PSII photodamage (Davis *et al*., 2016). In addition, we found increased sensitivity to ROS, as well as increased superoxide tissue staining in the Δ*gr1* mutant. The chloroplasts possess a very efficient removal system for ROS, with several ascorbate peroxidases that detoxify hydrogen peroxide using ascorbate as an electron donor. On the other hand, ascorbate peroxidases represent themselves prominent targets for ROS-induced damage (Dietz, 2016). Therefore, it is possible that the higher ROS fluxes reached in the Δ*gr1* plants, even under non-stress conditions, leads to decreased enzymatic ascorbate peroxidation and increased ROS-induced damage. Further, ROS inhibit plastid translation and thereby PSII repair (Nishiyama *et al*., 2011). In tobacco, 30 % of plastid GR activity was sufficient to avoid ROS formation under non-stress conditions (Ding *et al*., 2009). Our data clearly indicate that the presence of functional PSII in increasing light intensities is linked to a highly reducing stromal *E*_GSH_.

Only a low percentage of quantified protein differed in protein abundance between Δ*gr1* and WT plants in LL and after a shift from LL to HL (7.5 %, 125 of 1657). The affected proteins are distributed across several compartments, with the plastid being the most prominent localisation (30 %), confirming the important role of GR for plastid processes. The impact also on cytosolic proteins shows that the mutant cells adjust their protein content to the altered situation in the chloroplasts, possibly suggesting active retrograde signalling between chloroplast and nucleus. Notably, after the shift to HL, also mitochondrial proteins (3 complex I subunits) became affected, suggesting a role of mitochondrial GR under HL conditions.

Following manual annotation of proteins and allocation to process categories several patterns became apparent. Proteins involved in cytosolic translation were mostly less abundant in Δ*gr1* plants, while several proteins from plastid translation were more abundant. In addition, proteins of plastid glycolysis and fatty acid biosynthesis as well as proteolysis- and protein folding-related proteins were more abundant in Δ*gr1* plants. Adjustment of several photosynthesis-related proteins was already apparent in LL, confirming that GR function is not only relevant under stress conditions.

In HL, altered levels of transport proteins such as the inner envelope H^+^/K^+^ antiporter (KEA) isoform may contribute to phenotypes such as decreased ∆pH within the *pmf* component in Δ*gr1* mutants (Kunz *et al*., 2014). Interestingly, plant KEA isoforms possess sequence similarity to the glutathione-regulated potassium-efflux system KefC of *Escherichia coli* (Roosild *et al*., 2010). This system is important for protection against toxic electrophiles via acidification (Ferguson, 1999) and is negatively regulated by GSH, but activated by glutathione conjugates (Roosild *et al*., 2010). It is tempting to speculate that activity or abundance of plant KEA isoforms in the inner plastid envelope are linked to changes in stromal glutathione redox state. In the absence of AtKEA1/2 NPQ decreased and PSII was compromised (Kunz *et al*., 2014), which is line with the Δ*gr1* data. In contrast, deletion of the thylakoid localized H^+^/K^+^ antiporter AtKEA3 resulted in high NPQ in Arabidopsis (Armbruster *et al*., 2014; Wang *et al*., 2017). As stated above, analyses of *pmf* partitioning between ΔΨ and ΔpH revealed a decrease in the putative ΔpH component. Notably, the ΔpH component could be also modulated by ATP hydrolysis/formation in the dark until equilibrium between the *pmf* and the phosphorylating potential is reached (Cruz *et al*., 2001; Allorent *et al*., 2018). Considering the less reducing stromal *E*_GSH_ in Δ*gr1*, a difference in phosphorylating potential between WT and Δ*gr1* is possible, which could also explain differences in the putative ΔpH component.

Notably, the abundance of several known redox-regulated proteins was decreased in absence of GR1, such as one FBPase isoform and the chloroplast ATP-synthase gamma subunit. It is tempting to speculate that cysteines may become over-oxidised in Δ*gr1* plants, leading to changes in protein activity or to protein degradation (De Smet *et al*., 2019; Zaffagnini *et al*., 2019).

In conclusion the overall phenotype of Δ*gr1* mutants is likely a mixture of different effects: direct consequences via *E*_GSH_ and protein mis-regulation, and indirect consequences via ROS-induced protein damage. Investigations of cysteine redox state, GSH metabolites, glutathionylation of proteins and redox-cascades downstream of plastid/mitochondrial *E*_GSH_ provide scope for future research.

## Acknowledgements

This work was funded in part by grants from the Deutsche Forschungsgemeinschaft (DFG) to A.J.M., S.J.M., M.Schw. and P.D. within the research training group RTG2064 (‘Water use efficiency and drought stress responses: From Arabidopsis to barley’), the priority programme SPP1710 ‘Dynamics of thiol-based redox switches in cellular physiology’ (grants ME1567/9-1, SCHW1719/7-1) and by the Excellence Initiative of the German Federal and State Governments (EXC 294 to R.R.). We are grateful for support via the “Plant Mitochondria in New Light Initiative” (DFG, PAK918) including the grants SCHW1719/5-1 and MU4137/1-1. M.H. acknowledges funding by the Deutsche Forschungsgemeinschaft (DFG, HI 739/13-1). Research in S.K.’s laboratory is funded by DFG (EXC1028).

We thank Bastian Welter (University of Cologne) and Alexa Brox (INRES, University of Bonn) for technical assistance.

## Author Contributions

S.J.M.S., P.D., M.Schw., R.R., M.H. and A.J.M. planned and designed the research. S.J.M.S., R.W., D.D.G., J.R., S.K., M.R., V.L. performed experiments. M.Scho. analysed data. S.J.M.S. and A.J.M. wrote the manuscript. All authors discussed data and approved the final version of the manuscript.

**The following Supporting Information is available for this article:**

Fig. S1 Identification of PpGR1 knock-out mutants.

Fig. S2 Δ*gr1* mutant phenotype details.

Fig. S3 The ultrastructure of chloroplasts is not disrupted in Δ*gr1* mutants.

Fig. S4 Δ*gr1* plants are viable in extended periods of darkness.

Fig. S5 Δ*gr1* plants cannot profit from higher light fluencies.

Fig. S6 Δ*gr1* plants are light-sensitive - measurements of CEF and Fv/Fm.

Fig. S7 Experimental setup for protein quantification via metabolic labelling. Fig. S8 Verification of GR1 absence in Δ*gr1* by targeted LC-MS/MS.

Table S1 Primer list.

Table S2 Proteomics data and functional annotation (separate file).

