## Supplemental Information for "Chloroplasts Require Glutathione Reductase to Balance Reactive Oxygen Species and Maintain Efficient Photosynthesis"

**The following Supporting Information is available for this article:**

**Fig. S1 Identification of PpGR1 knock-out mutants.**

**Fig. S2 Δ*gr1* mutant phenotype details.**

**Fig. S3 The ultrastructure of chloroplasts is not disrupted in Δ*gr1* mutants.**

**Fig. S4 Δ*gr1* plants are viable in extended periods of darkness.**

**Fig. S5 Δ*gr1* plants cannot profit from higher light fluencies.**

**Fig. S6 Δ*gr1* plants are light-sensitive - measurements of CEF and Fv/Fm.**

**Fig. S7 Experimental setup for protein quantification via metabolic labelling.**

**Fig. S8 Verification of GR1 absence in Δ*gr1* lines by targeted LC-MS/MS.**

**Table S1 Primer list.**

**Table S2 Proteomics data and functional annotation (separate file)**

**Methods S1+Additional References**

**Fig. S1 Identification of PpGR1 knock-out mutants.**

**(a)** Gene model of *PpGR1* showing the intron and exon structure, as well as the regions chosen for homologous recombination (5' HR, 3' HR). A large part of the coding sequence including the active site (arrow) was replaced by a hygromycin resistance cassette (*hpt*).
**(b)** In regenerated plants that survived Hygromycin selection, integration of the construct into the target locus was verified using primer pairs spanning the 5' integration site (5P_F and H3b_R, Table S1) and the 3' integration site (NosT_F and 3P_R, Table S1).
**(c)** Absence of *PpGR1* transcript for knock-out lines #48 and #88 was confirmed using the primer pair PpGR1_RT_F and PpGR1_RT_R (Table S1). As a control for the presence of cDNA, *PpEF1alpha* transcript was used as control (Table S1, EF1a_RT_F and EF1a_RT_R2).

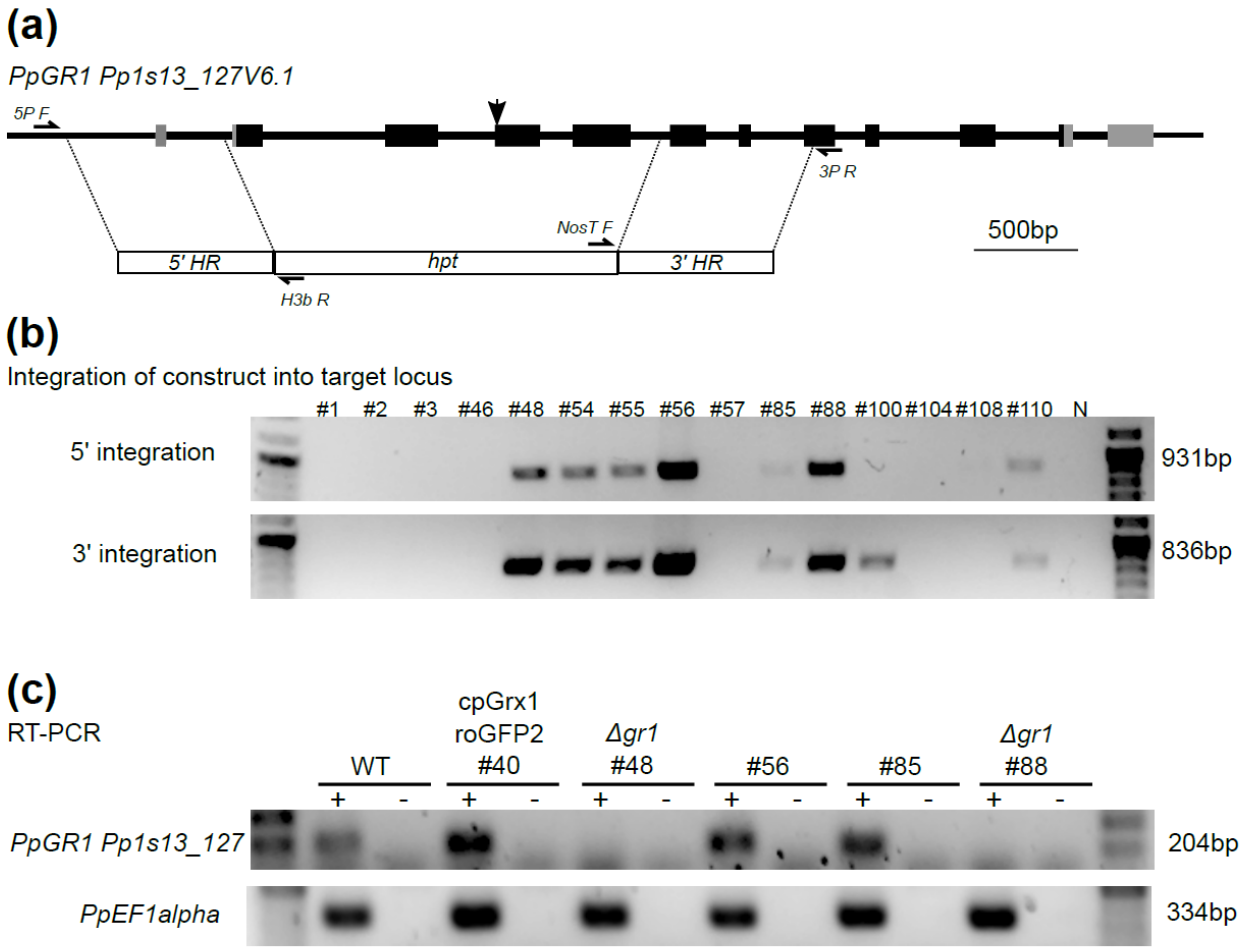

**Fig. S2 Δ*gr1* mutant phenotype details.**

**(a)** Phenotypic appearance of a line with plastid-localised Grx1-roGFP2 and two independent GR1 knock-out mutants after 23 d regeneration from protoplasts in a 16 h light/ 8 h dark regime. Growth of Δ*gr1* mutants is slow, colony diameter is smaller than in WT background. Bars, 1 mm.

**(b)** Δ*gr1* mutants do form less caulonema (arrowhead). Bars, 0.5 mm.

**(c)** Δ*gr1* mutants do form less rhizoids (arrowheads) at the base of gametophores. Bars, 0.1 mm.

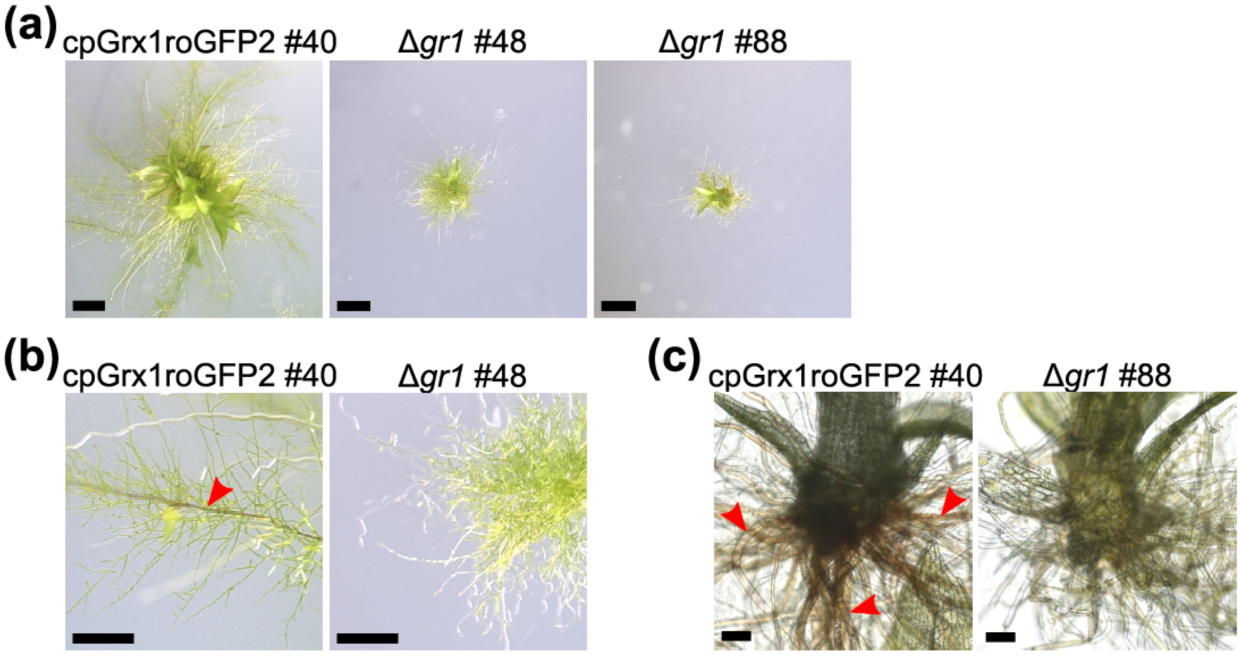

**Fig. S3 The ultrastructure of chloroplasts is not disrupted in Δ*gr1* mutants.**

**(a)** Transmission electron micrograph (TEM) of chloroplasts in protonema from liquid cultures. Bars, 0.5 µm.
**(b)** TEM of chloroplasts in gametophores from plate culture. Bars, 0.5 µm.

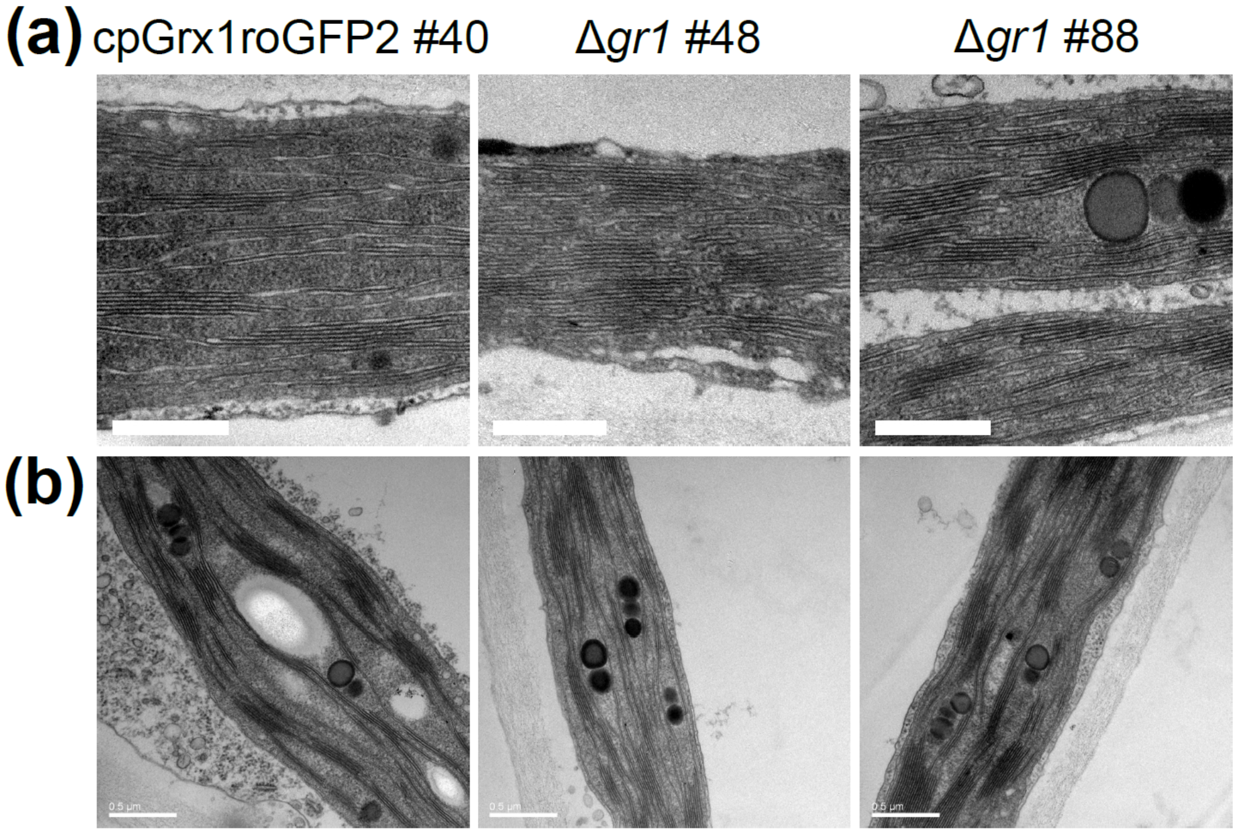

**Fig. S4 Δ*gr1* plants are viable in extended periods of darkness.**
**(a)** Phenotypic appearance of WT, a line with chloroplast-localised Grx1-roGFP2 and two independent GR1 knock-out mutants (Δ*gr1)* under different light regimes. CL, continuous light; SD, short day. Growth is restricted in the shorter light period, but mutants are viable. Bars, 1 mm.
**(b)** Δ*gr1* plants survive an extended incubation in darkness and fully recover, comparable to the sensor line in WT background. Bars, 1mm.

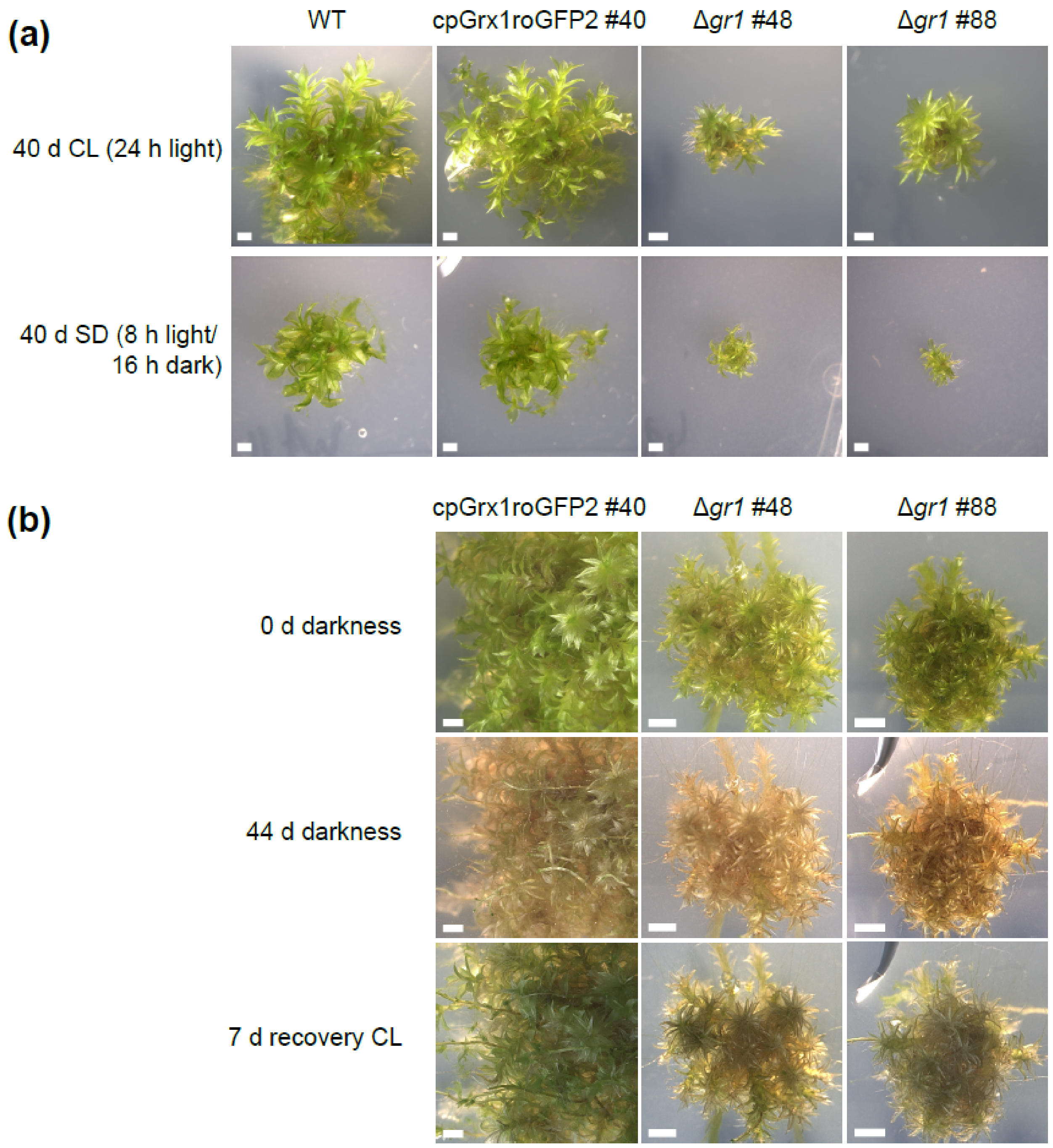

**Fig. S5 Δ*gr1* mutants cannot profit from higher light fluencies.**

**(a)** Phenotypic appearance of WT, a line with chloroplast-localised Grx1-roGFP2 and two independent GR1 knock-out mutants (Δ*gr1*) under different light intensities in a 16 h light: 8 h dark regime. Growth of Δ*gr1* mutants is not increasing with increasing light fluencies. Bars, 1 mm.

**(b)** Fresh weight of moss colonies under different growth light intensities. Growth of Δ*gr1* mutants is significantly different from WT lines in all tested light intensities ***P<0.001 (two-way ANOVA, TukeyHSD post hoc test). Growth is not increasing with increasing light fluencies, in contrast to moss lines with WT background. The plot depicts *n*=156 observations of colony fresh weight of GR1 ko lines (Δ*gr1*) and lines with GR1 wildtype background (WT). Plot: Line = mean, box = standard deviation, dots = data points.

**(a)**

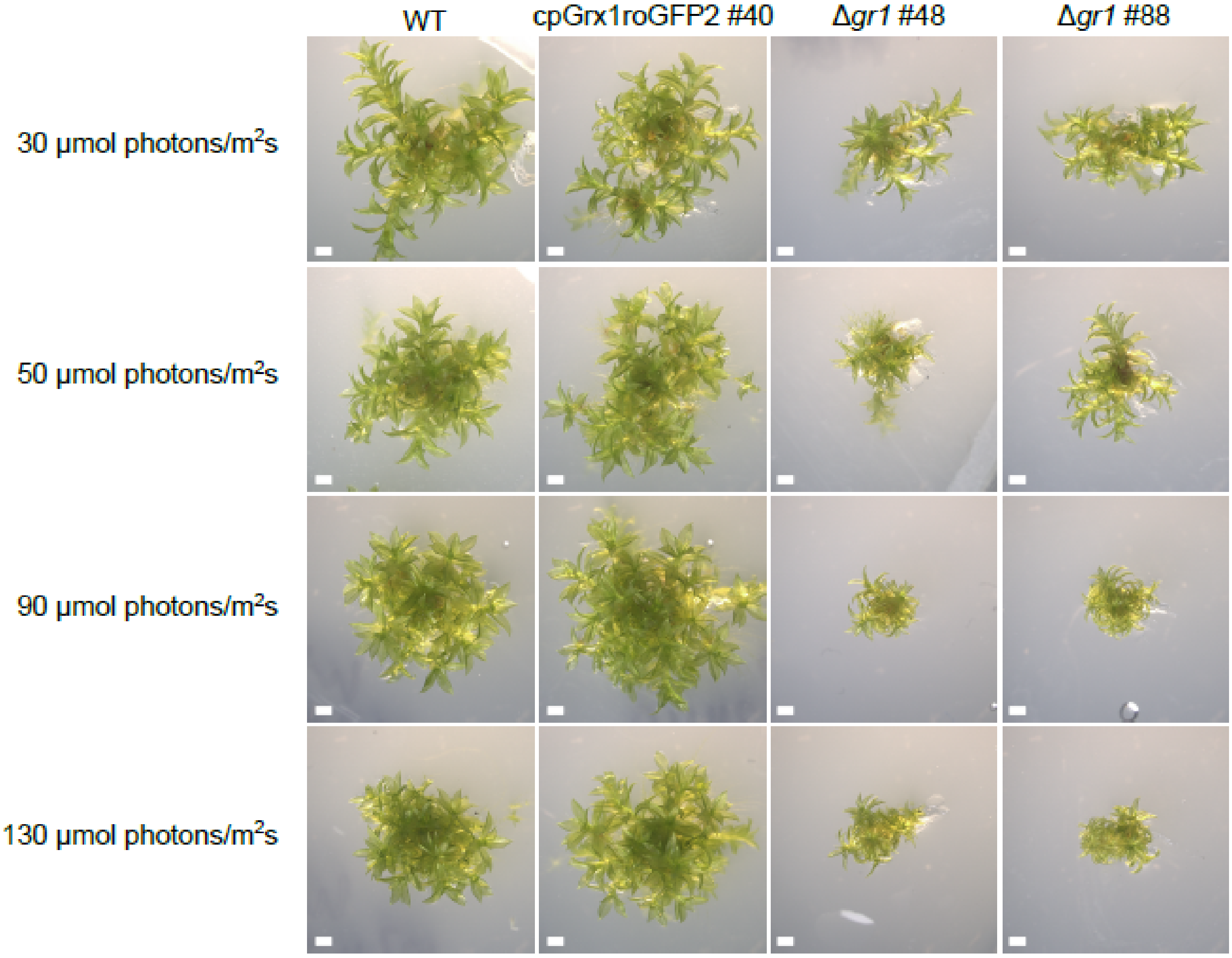
WT
cpGrx1roGFP2 #40
Δgr1 #48
Δgr1 #88
30 µmol photons/m²s
50 µmol photons/m²s
90 µmol photons/m²s
130 µmol photons/m²s

**(b)**

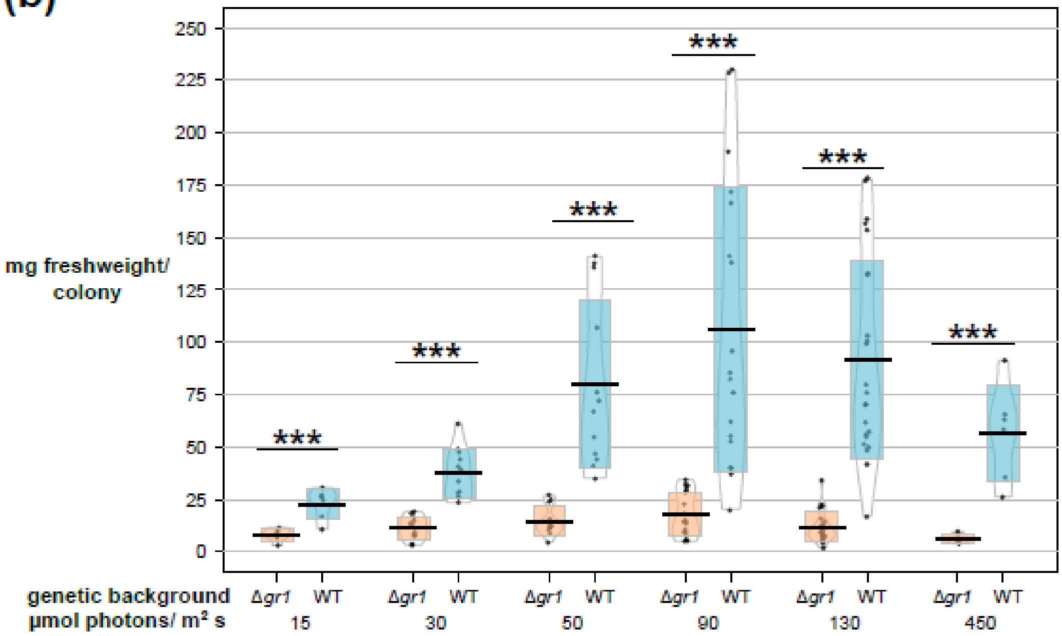
250
225
200
175
150
125
100
75
50
25
0
mg freshweight/ colony
***
genetic background
µmol photons/ m² s
Δgr1
WT
15
30
50
90
130
450

**Fig. S6 Δ*gr1* mutants are light-sensitive - measurements of CEF and Fv/Fm.**

**(a)** Photosynthetic parameters measured under low light (LL, 15 µmol photons $m^{-2}$ $s^{-1}$), control light (C, 50 µmol photons $m^{-2}$ $s^{-1}$) and high light (HL, 450 µmol photons $m^{-2}$ $s^{-1}$). Cyclic electron flow (CEF) is not significantly altered in medium to high fluencies in Δ*gr1* plants (left panel). Fv/Fm ratio decreases in Δ*gr1* plants with increasing light, indicating decreased photosynthetic efficiency (right panel).

**(b)** Cyclic electron flow induced by anoxia is increased in Δ*gr1* plants after 6 h, $P<0.05$ ($n=6$, paired t-test).

**(c)** Photometric determination of chlorophyll (a+b) levels (gametophores, LL, 15 µmol photons $m^{-2}$ $s^{-1}$. $n=3$, small letters depict significant differences, $P<0.05$ (one-way ANOVA, Tukey HSD post hoc test).

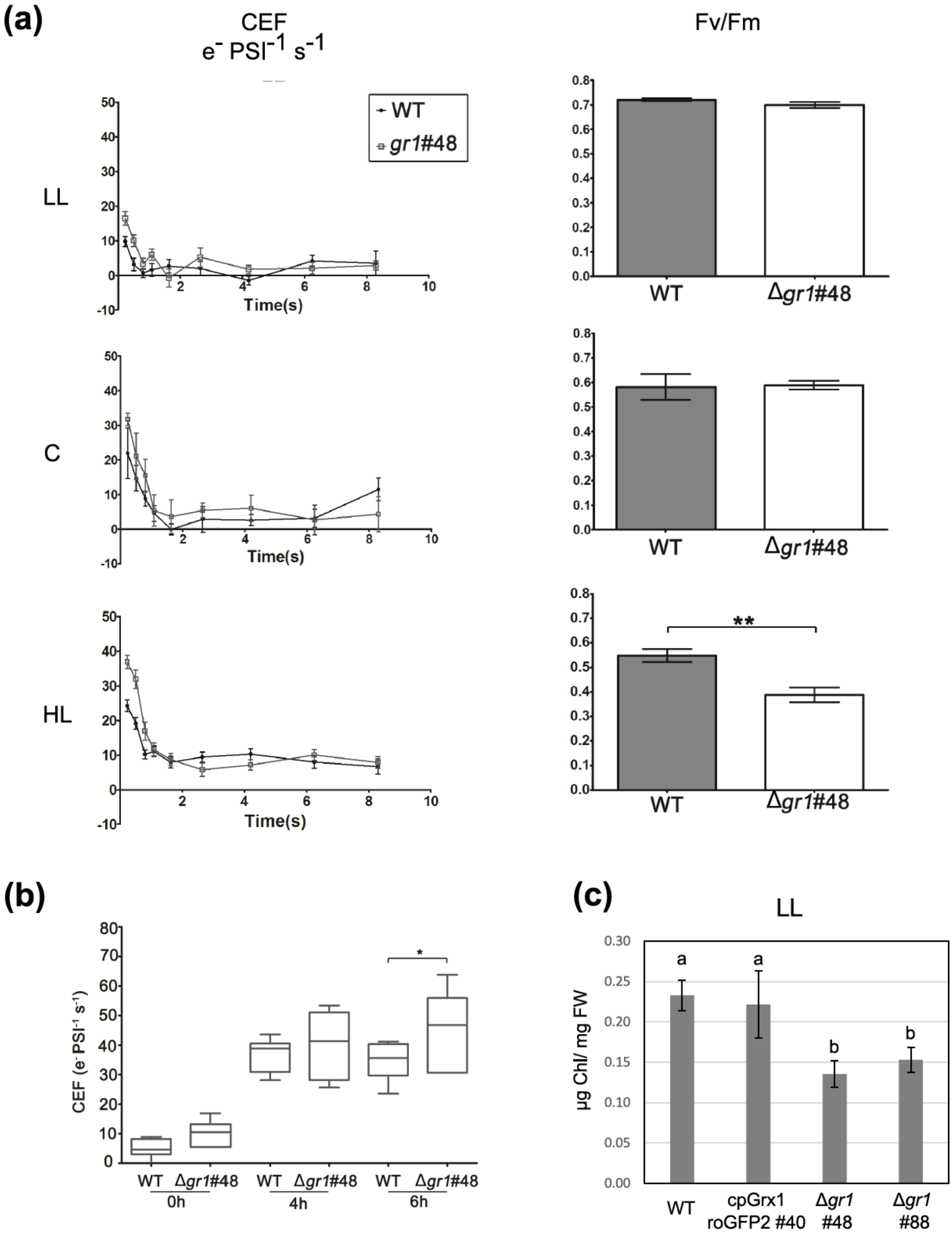
(a)
CEF
e⁻ PSI⁻¹ s⁻¹
Fv/Fm
LL
C
HL
WT
gr1#48
Time(s)
Δgr1#48
**
(b)
CEF (e⁻PSI⁻¹ s⁻¹)
WT
Δgr1#48
0h
4h
6h
*
(c)
LL
µg Chl/ mg FW
a
b
WT
cpGrx1 roGFP2 #40
Δgr1 #48
Δgr1 #88

**Fig. S7 Experimental setup for protein quantification via metabolic labelling.**
**(a)** *Physcomitrella patens* protonema was grown in low light and transferred to high light for 1 h. In the experimental replicates, the heavy nitrogen label ($^{15}N$) was swapped between the cpGrx1roGFP2 #40 line (WT background) and Δ*gr1 #48*. Protein extracts from WT samples and Δ*gr1* were mixed 1:1 to compare protein abundances.

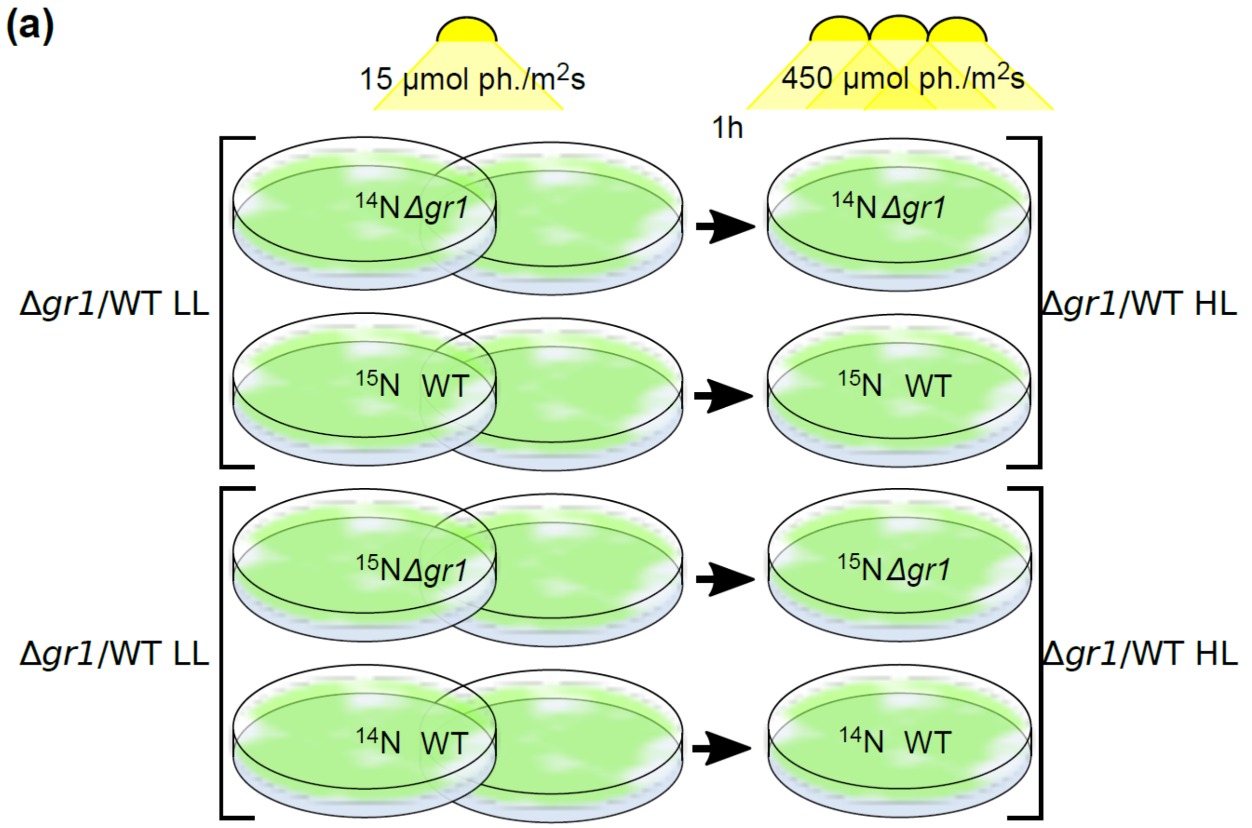

**Fig. S8: Verification of GR1 absence in Δ*gr1* lines by targeted LC-MS/MS.**
Total protein from wildtype (WT) and two independent GR1 knock-out lines (Δ*gr1* #48, Δ*gr1* #88) was digested with trypsin and GR1 peptides were monitored by targeted LC-MS/MS (PRM). Peptides derived from ATPase subunit beta (ATPB) served as loading controls. GR1 peptides (in order of elution): IIVDASSDK, TAFGNEPTKPDYR, ATVPNIPGK, SEVDLIEGR, NLGLEEVGVK, GFGWSFETEPK, EYDYDLIAIGAGSGGVR, TNVDSIWAIGDVTNR. ATPB peptides (in order of elution): VVLEVAQHLGENTVR, VVDLLAPYQR, FTQANSEVSALLGR, IINVIGEAIDER.

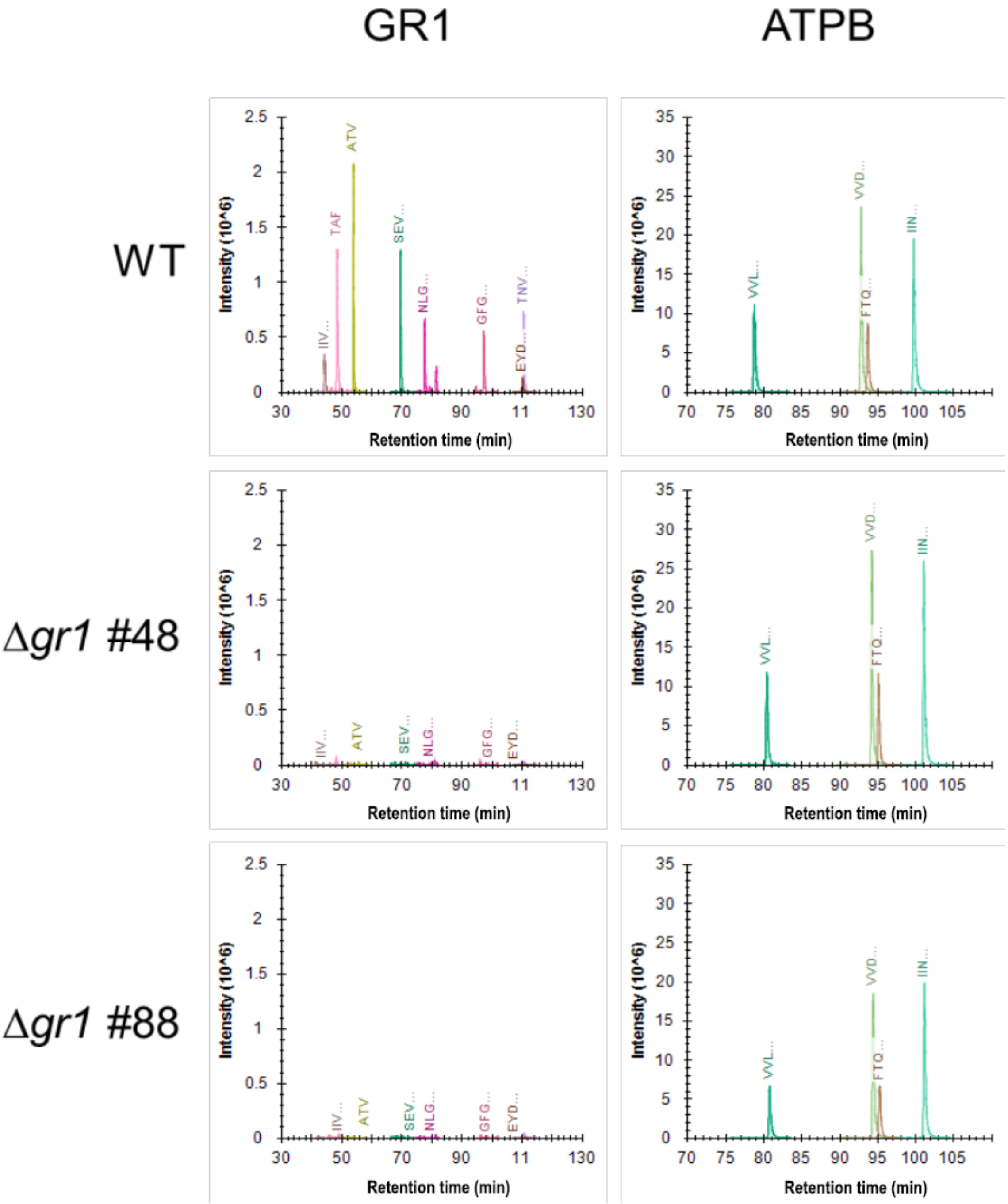

**Table S1** Primer list.

| Primer name | Sequence (5'->3') | Purpose |
|---|---|---|
| PpGR1ko_5PHR_F | gctcttcaaaacaggatttgatagcc catc | Amplify 5' homologous region |
| PpGR1ko_5PHR_R | cagatccttggcggcccgaatccca atgtgctagg | Amplify 5' homologous region |
| PpGR1ko_3PHR_F | tatgttactagatcgtcttgagggatgt tctgcga | Amplify 3' homologous region |
| PpGR1ko_3PHR_R | gctcttcaatgcggtttgtcacgtctc | Amplify 3' homologous region |
| PpGR1ko_hpt_F | cacattgggattcggGCCGCCA AGGATCTGATG | Amplify hpt cassette |
| PpGR1ko_hpt_R | gaacatccctcaagaCGATCTA GTAACATAGATGACACC G | Amplify hpt cassette |
| 5P_F | gaagcacaacaaagagaggca | Detect 5' integration GR1 ko construct |
| H3b_R | CCAAACGTAAAACGGCT TGT | Detect 5' integration GR1 ko construct |
| NosT_F | gcgcggtgtcatctatgtta | Detect 3' integration GR1 ko construct |
| 3P_R | ggctcatttccgaaagcagt | Detect 3' integration GR1 ko construct |
| EF1a_RT_F | cgacgcccctggacatc | Amplify EF1alpha transcript |
| EF1a_RT_R2 | CATGTTGTCACCCTCGAA CC | Amplify EF1alpha transcript |
| PpGR1_RT_F | CTATCGGGGCTGGTAGT GG | Amplify GR1 transcript |
| PpGR1_RT_R | GGCTGCTCTCAAAATCG TGT | Amplify GR1 transcript |

**Methods S1**

**Anoxia-induced CEF**

To measure cyclic electron flow under anoxia, protonemal tissues were homogenised and resuspended in $PpNO_3$ medium and incubated for 6 days under LL before treatment with argon bubbling. For photosynthetic electron transfer measurements, samples were taken after 0 h, 4 h and 6 h.

**Pigment analysis: Chlorophyll content**

Plant tissue was frozen in liquid nitrogen, homogenised using a Retsch mill and extracted with 80% acetone followed by a second extraction with 100% acetone. Combined supernatants were measured with a solvent gradient (solvent A acetonitrile/methanol/0.1 M Tris-HCl (72:8:3, (v/v)) and solvent B methanol/ethyl acetate (68:32, (v/v)) by HPLC according to Thayer & Björkman (1990). The amount of chlorophyll per mg fresh weight in gametophore samples was determined from the same acetone extracts as used for HPLC analysis (Thayer & Björkman, 1990) by photometric measurement according to Porra *et al.* (1989).

**Verification of GR1 knock-out by Parallel Reaction Monitoring (PRM)**

Hardware configuration of the chromatography system and eluents were the same as described above. The gradient for peptide separation was programmed as follows: 2.5-18 % (v/v) B over 55 min, 18-32 % (v/v) B over 50 min, 32-99 % (v/v) B over 5 min, 99 % (v/v) B for 20 min. The Q Exactive Plus was operated with the following PRM settings: resolution: 70,000 at m/z 200, AGC target: 5e4, maximum injection time: 200 ms, isolation window: 1.6 *m/z*. A target list containing several tryptic GR1 peptides, identified by previous analyses in our lab, was compiled with Skyline (version 4.1; (MacLean *et al.*, 2010)) and used for scheduled fragmentation of GR1 peptides. In addition, four peptides of ATPase subunit beta (Pp1s310_30V6.2), which served as loading controls, were included in the target list. After data acquisition MS raw files were loaded into Skyline for peak extraction. The presence of minimum of four fragment ion traces per peptide with a dotp of at least 0.95 was regarded as safe peptide identification.
